# ExSEnt for explainable dementia detection: disentangling temporal and amplitude-driven complexity boosts EEG-based classification

**DOI:** 10.64898/2026.02.02.703144

**Authors:** Sara Kamali, Fabiano Baroni, Pablo Varona

## Abstract

Early detection of dementia enables timely intervention and better care planning. Electroencephalography, being accessible and noninvasive, offers a practical avenue for monitoring pathological alterations in neural activity. Classical biomarkers like the theta-to-alpha power ratio (TAR) along with complexity measures are common methods that are usually evaluated and used for dementia detection. In this study, we aimed to assess the discriminative ability of a novel entropy-based family of measures, Extrema-Segmented Entropy (ExSEnt), summarized by multiple robust statistics per subject for dementia, along with classical measures, and evaluate the incremental value of these metrics. We analyzed an EEG dataset comprising healthy controls, individuals with Alzheimer’s disease, or frontotemporal dementia. Following preprocessing and group-level analyses of independent components, we focused on source-space activity from the prefrontal cortex and visual association cortices—regions implicated in early disease. From these sources, we computed complexity metrics: Sample Entropy, Katz Fractal Dimension, Higuchi Fractal Dimension, and Hurst exponent and ExSEnt metrics along with TAR and band-limited power at delta, beta, low and high gamma bands. Using stability-based selection with elastic net logistic models, we identified a reliable set of discriminative features and quantified their cross-subject robustness. This framework isolates interpretable and trustworthy source-local biomarkers from single-region time series. We observed that the alpha/theta temporal entropy measures (ExSEnt) are selected as the most reliably informative metrics in the left prefrontal cortex, yielding a classification performance comparable to what was recently reported with high-dimensional deep learning methods for this dataset, with a simple logistic regression model on a single brain source.

## 1. Introduction

Dementia is characterized by cognitive decline severe enough to interfere with daily functioning. While the term refers to the syndromes (memory loss, executive dysfunction, language, and other cognitive domains), the underlying causes can be varied [1, 2, 3]. The estimated prevalent cases of dementia by the year 2050 are around 152 million people worldwide [4]. The most common form is Alzheimer’s Disease (AD) (accounting for roughly 60–80% of cases [5]) and Frontotemporal Dementia (FTD), also called frontotemporal lobar degeneration, which accounts for about 10% of all dementia cases [6].

EEG provides a noninvasive and cost-effective measure of brain activity with high temporal resolution, making it suitable for recording and monitoring neural activity. Many EEG-based dementia studies rely on feature extraction followed by statistical or machine-learning classification. In AD, restingstate EEG shows a robust slowing of background rhythms [7, 8, 9, 10, 11], commonly summarized by an increased theta-to-alpha power ratio (TAR) [12, 13]. FTD may also show slowing, but it is often milder in early stages, with limited frontal slowing [14, 15, 16, 17], and a higher dominant peak frequency compared with AD [18]. In AD, these spectral changes track cognitive impairment and disease severity [8, 10].

Complexity metrics capture nonlinear, multiscale properties of brain dynamics and have become effective features for ML classification of dementia. In AD and along the AD continuum, numerous studies report reduced signal irregularity, together with ‘decreased fractal dimension (e.g., Higuchi/Katz) and Lempel–Ziv complexity, consistent with diminished dynamical richness; these alterations are further evident in microstate sequence complexity and in network-level entropy [19, 20, 21, 22, 23, 24, 25, 26, 27, 28]. Largest Lyapunov exponent analyses indicate reduced chaoticity of cortical dynamics with aging and cognitive decline and have been proposed as complementary biomarkers within AD EEG pipelines [29]. When fed into classifiers such as SVMs, random forests, or deep networks (e.g., CNNs, LSTMs), EEG complexity features, alone or combined with spectral and connectivity descriptors, have shown robust discrimination of AD, MCI, and HC, and more recently have also supported differentiation of AD from FTD [30, 31, 32, 33]. Methodological advances, e.g., robust multiscale fuzzy entropy for EEG [21], fractal-dimension distributions that capture spectral–scale interactions [26], and network-level permutation entropy in M/EEG [34], further improve generalization, supporting complexity-informed, low-cost pipelines for early dementia screening and subtype classification.

ExSEnt is a novel complexity feature family designed to increase sensitivity to both temporal and amplitude-driven irregularities in time series by defining seperate entropy measures for each, making it a plausible biomarker for distinguishing pathological from healthy brain dynamics [35]. Previous works have shown that temporal irregularity is a promising biomarker of AD [36]. Here, we evaluated classification using a reference feature set of established spectral and nonlinear EEG descriptors and quantified the incremental predictive value of adding ExSEnt.

To transition from high-dimensional complexity metrics to a clinically viable biomarker pipeline, we performed feature selection to identify a parsimonious and informative feature subset. In biomedical data analysis, feature selection methods are generally categorized into three types: filter, wrapper, or embedded approaches [37, 38, 39]. Filter methods, such as univariate statistical tests, mutual information, or correlation-based ranking, evaluate each feature independently of a classifier, efficiently filtering out irrelevant variables [40]. Wrapper methods, including recursive feature elimination or sequential forward and backward selection, iteratively evaluate candidate subsets by repeatedly fitting the downstream model, optimizing predictive performance at the cost of higher computational complexity [41, 42]. Embedded methods, such as LASSO, elastic net, or tree-based importance measures, incorporate feature selection directly into model training through regularization or internal weighting mechanisms [43, 44].

While each approach has merits, a fundamental limitation of these methods, particularly in high-dimensional, low-sample-size settings common in EEG analysis, is their instability. Small perturbations in the data, such as differences in resampling, noise, or cross-validation splits, can lead to substantially different selected features. This sensitivity severely undermines the reproducibility and biological interpretability of the results, casting doubt on whether identified biomarkers reflect true pathophysiology or are merely statistical artifacts [45, 46, 47].

To address this challenge, we adopt stability selection, a robustness-oriented hybrid method originally proposed by Meinshausen and Bühlmann [48] and later refined in numerous studies [49, 50]. Stability selection combines subsampling with penalized models, such as the elastic net, to repeatedly fit the data on random subsets of samples and estimate how consistently each feature is selected across iterations. Features exceeding a predefined selection-frequency threshold are retained as stable predictors. This framework offers several advantages for biomarker discovery: it effectively controls false-positive inclusion through theoretical error bounds on the expected number of false selections [48], emphasizes reproducibility by prioritizing features that generalize across subsamples [47, 45], mitigates overfitting in high-dimensional and small-sample settings common in EEG analyses, and remains model-agnostic, allowing integration with various base learners.

Advancement of machine learning (ML) and deep learning (DL) has enabled automated analysis of complex neuroimaging and electrophysiological data [51, 52]. Large-scale structural MRI, MEG, and EEG datasets have been employed to train classifiers that distinguish AD, FTD, and healthy controls with increasing accuracy [53, 54, 55, 56]. Classical ML models, such as support-vector machines (SVMs) and *k*-nearest neighbors (*k*-NN), applied to spectral power and connectivity features of EEG, have demonstrated reliable differentiation of dementia from controls [57, 58]. More recently, deep learning models, including CNNs and recurrent neural networks (RNNs), have shown the ability to extract spatiotemporal EEG features and microstate dynamics, further enhancing classification performance [59, 60, 61]. In sum, ML and DL methods are rapidly improving early-dementia screening by integrating imaging, EEG, genetics, and clinical data into powerful predictive models, with the potential to support precision diagnostics and targeted interventions [62, 63].

The rise of explainable artificial intelligence (XAI) has added interpretability to otherwise opaque predictive models. Rather than functioning as “black box” classifiers, techniques such as SHAP (SHapley Additive exPlanations) and LIME (Local Interpretable Model-agnostic Explanations) reveal which features drive individual decisions [64, 65]. Applied to EEG, XAI has highlighted posterior slowing in AD [66], frontal–temporal sources distinguishing AD from FTD [67], and temporal theta and parietal alpha rhythms as key markers [68]. Such transparency helps validate AI-derived biomarkers against known neuropathology and increases clinical trust in automated tools, facilitating their translation into clinical practice [69, 70].

Explainable methods boost reliable early diagnosis of dementia, which is critically important because identifying individuals in the mild cognitive impairment (MCI) stage or prodromal dementia can dramatically affect management and outcomes [71, 72] and delay cognitive decline [73, 74]. Moreover, characterization of early neural biomarkers is essential for drug development, since disease-modifying therapies for dementia may only be effective if given during preclinical or prodromal stages [75, 76].

In this work, we used an EEG dataset to quantify the stability of complexity-based measures and evaluated their reliability as biomarkers for dementia detection. In our pipeline, we applied stability selection over spectral and complexity-based EEG features, including ExSEnt metrics. Only features surpassing a reproducibility threshold were retained for subsequent logistic regression (LR). This ensured that downstream analyses rely on features that were not only predictive but also robust and interpretable, forming a reliable foundation for explainable biomarker discovery in dementia.

Explainability in this setting arises from the information disentangled inputs, the model class, and from the selection procedure: LR provides direct, signed coefficients, while stability selection yields selection probabilities that quantify how consistently each feature is retained under resampling. Together, this approach provides explainability by design, because the model decision is anchored to a small set of reproducible features with explicit directions and magnitudes of association with pathology, rather than post-hoc explanations.

In the following sections, we introduced the dataset and the preprocessing method for group-level brain source feature extraction. Then, to isolate the effect of ExSEnt, we compared two otherwise identical stability selection pipelines: one trained without ExSEnt (WoE) and one trained with ExSEnt added to the same reference feature set (WE). Both pipelines used the same preprocessing, scaling, cross-validation strategy, and model class; ExSEnt was the only added information in WE. Performance was reported separately for each region. We interpret consistent performance gains together with the stability of ExSEnt feature selection across resampling.

## 2. Method

### 2.1. Dataset

We utilized a publicly available resting-state, eyes-closed EEG dataset encoded in BIDS (Brain Imaging Data Structure) format [77]. The dataset comprises 88 subjects: 36 with AD, 23 with FTD, and 29 age-matched healthy controls (HC). Participants’ mean ages were 66.4±7.9 years in the AD group, 63.7±8.2 years in the FTD group, and 67.9±5.4 years in the HC group, indicating comparable age distributions across groups. Male–female ratios differed: the AD group included relatively more female participants, whereas the FTD and control groups had a higher male representation (Table 1). EEG was recorded using a standard clinical 19-channel monopolar montage, with a sampling rate of 500 Hz and an average recording duration of approximately 13 minutes per subject (ranging from 5 to 21 min). Both raw and rigorously preprocessed EEG data—cleaned via Artifact Subspace Reconstruction (ASR) and Independent Component Analysis (ICA)—are provided online [77]. This study was conducted on the raw EEG data, to which we applied a preprocessing pipeline tailored to the objectives of our analysis. Demographic and clinical metadata include Mini-Mental State Examination (MMSE) scores, widely used for cognitive screening in dementia research [78]. This well-characterized dataset supports machine learning and nonlinear signal analysis aimed at discriminating AD, FTD, and healthy aging using both spectral and complexity features.

**Table 1:**
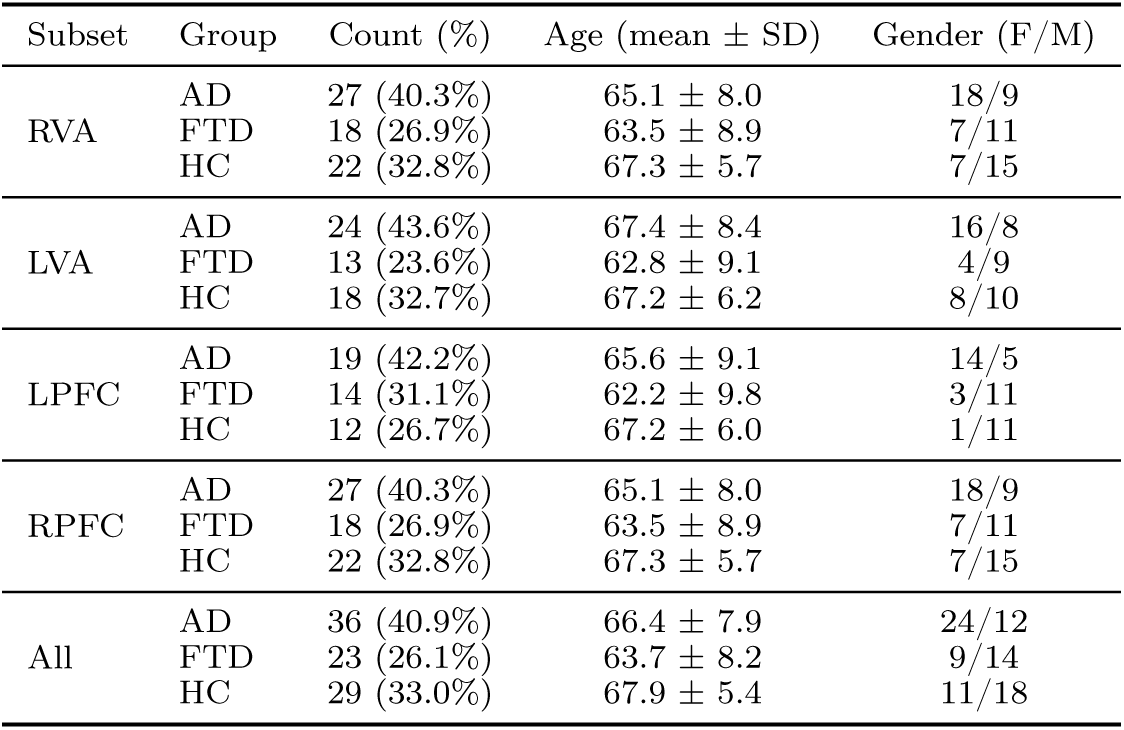
Demographic summary of subjects in all clusters.

### 2.2. Data preprocessing

We applied a comprehensive preprocessing pipeline to extract brain sources and enable group-level analysis. First, continuous EEG data were bandpass filtered from 1 to 100 Hz using a zero-phase FIR filter implemented via MATLAB’s filtfilt, preserving phase integrity. Line noise at approximately 50 Hz and its harmonic at 62.5 Hz were attenuated using the CleanLine EEGLAB plugin, which adaptively estimates and removes sinusoidal noise [79]. ASR was then applied to reconstruct corrupted data segments and remove noisy channels, via EEGLAB’s clean-rawdata plugin [80]. After average referencing, we decomposed the cleaned data into Independent Components (ICs) using Adaptive Mixture Independent Component Analysis (AMICA), an efficient ICA method offering robust source separation for EEG [81, 82]. Equivalent dipole modeling was performed using EEGLAB’s DIPFIT plugin, configured under the MNI boundary-element head model for source localization, with a coarse-to-fine grid search to optimize dipole fitting. Electrode positions were co-registered to the head model, and resulting dipoles were assigned to Talairach brain regions. Only ICs exhibiting residual variance (RV) ≤ 15% and classified as brain sources with ≥ 60% probability (via EEGLAB’s automatic ICLabel) were retained. Ocular artifacts were also saved to be used for clustering sanity checks. A schematic of a similar preprocessing workflow—with specific adaptations—is available in [83], Fig. 2.

### 2.3. Group level clustering

Our objective was to identify comparable brain sources across participants, in order to perform classification and analysis at the group level within similar components. Hence, we applied clustering to the individually extracted ICs, grouping components with similar spatial characteristics to enable meaningful group-level comparisons. We used EEGLAB’s STUDY framework to manage and analyze ICs at the group level systematically. We included only the brain ICs deemed acceptable during preprocessing, along with eye components to verify the clustering accuracy (as these are expected to cluster together). Clustering was implemented using the *k*-means algorithm, with dipole location and scalp-map weights set to 10 and 1, respectively, to emphasize spatial source consistency. Since power was one of our target features for classification, we avoided using the power spectrum as a clustering metric. We specified *k* = 10 clusters (see Fig. 1). The number of clusters was chosen empirically by trial and error: we tested a range of *k* values and selected one at which the clustering became stable across repeated runs (i.e., the same configuration reappeared). As an additional check, we retained all eye-related ICs; if the majority consistently fell into the same cluster, this further supported that the chosen *k* was appropriate. If a cluster contained multiple ICs from the same participant (as all of them did), we retained only the component that was a better match to that cluster by taking into account the scalp topography and dipole location and comparing them with the group’s average. This prevented any single subject from disproportionately influencing a cluster. After this step, eight clusters were retained, each comprising ICs from more than 50% of the subjects in the dataset. We focused our subsequent analysis on four of the clusters (Fig. 2), localized to the right and left visual association cortices (RVA, LVA) and the right and left prefrontal cortices (RPFC, LPFC), as these areas are particularly implicated in dementia-related changes, as discussed in section 1.

**Figure 1:**
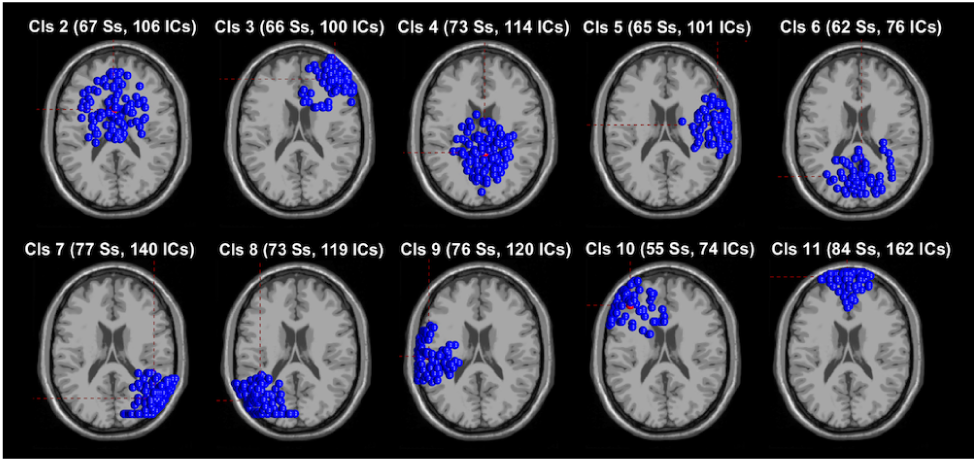
Initial ICA clustering across subjects, including both brain and ocular components. Cluster 11 contains only eye-related ICs. Subjects can contribute multiple ICs to a single cluster—the number of components exceeds the number of subjects in every cluster.

**Figure 2:**
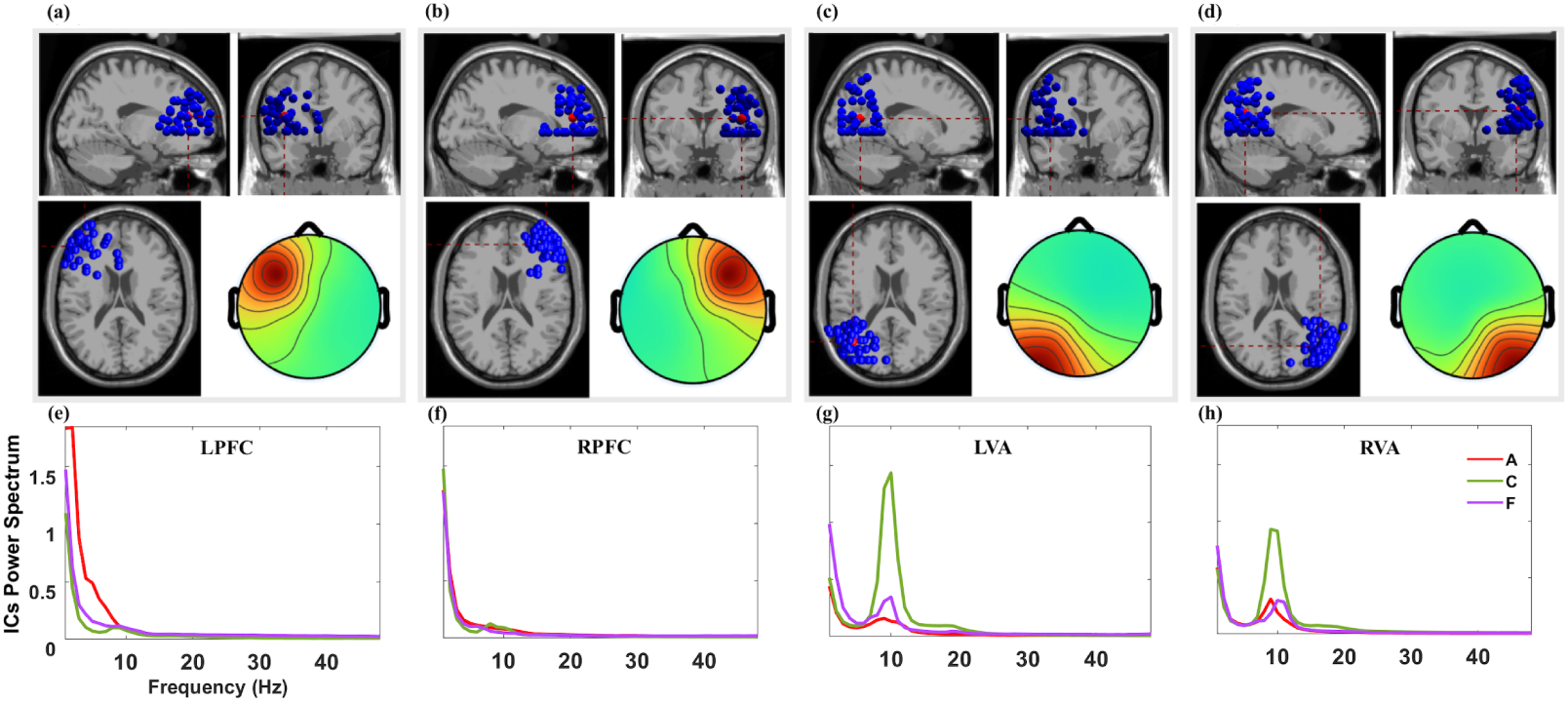
Final IC clusters selected for group-level analysis. Panels show: (a) left prefrontal cortex (LPFC) with 45 ICs; (b) right prefrontal cortex (RPFC) with 58 ICs; (c) left visual association cortex (LVA) with 55 ICs; and (d) right visual association cortex (RVA) with 67 ICs. Each subpanel displays sagittal (top left), coronal (top right), and axial (bottom left) views of the brain with the spatial locations of all ICs in that cluster. Each subject contributes at most one IC per cluster. The bottom right inset in the top row of panels is the average scalp map of the included ICs. Panels (e) to (h) show the average power spectrum of each group in the corresponding cluster, where red, purple, and green show the AD, FTD, and HC groups, respectively.

The bottom panels in Fig. 2 show the average power spectrum of each group within the corresponding brain area. As illustrated, the distinction between dementia and healthy groups was more pronounced in the left hemisphere. In the LPFC, the AD group exhibited higher delta and theta power than the other groups, while the FTD group showed slightly higher theta power than HC. In the contralateral cluster, RPFC, the differences among the three groups were only marginal. In the LVA, the FTD group showed higher average delta power than the other two groups, while in the alpha band, the HC group exhibited notably higher power, with AD displaying a lower peak than FTD. In the beta band, HC had slightly higher power than both dementia groups. In the RVA, FTD’s delta power was only marginally higher than the other groups. HC still showed higher alpha power in this cluster, although the difference relative to the dementia groups was reduced by ∼ 35% compared to the LVA. In the beta band, HC again showed slightly higher power than the other groups.

Table 1 summarizes the demographic distribution of subjects across the four selected clusters, along with all the subjects in the dataset. Each brain area contained comparable proportions of AD (40.3–43.6%), FTD (23.6–31.1%), and HC (26.7–32.8%) participants. The mean ages across groups were in the mid-60s, with standard deviations of approximately 6–10 years, and no systematic differences were observed between clusters. Gender composition showed a higher female-to-male ratio in AD groups for all the clusters, whereas HC and FTD groups showed a higher prevalence of male participants (particularly in the LPFC cluster), as observed in the total dataset.

### 2.4. Feature extraction

We applied a continuous wavelet transform (CWT) using analytic Morse wavelets to filter the components’ activity within the delta (–4 Hz), theta (4–8 Hz), alpha (8–12 Hz), beta (12–30 Hz), low gamma (30–48 Hz), and high gamma (52–100 Hz) bands. Then, the wavelet coefficients were tapered with a Kaiser window to smoothly attenuate edge frequencies before reconstructing the band-limited signal via inverse CWT. After discarding the first and last 3.5 s of each filtered IC trace to remove edge artifacts, we extracted the features over 24-second-long windows with 50% overlap. Since our target was comparisons and classification of the subjects, the length of the data was bounded by the shortest recording; hence, we extracted 23 consecutive windows per subject. From each window the following features were extracted:

- The ratio of the average power in the theta band to that in the alpha band (theta/alpha power ratio, TAR), together with the average band-limited power, across delta, beta, low and high gamma bands. Band-limited power is EEG biomarkers in dementia [84], reflecting the characteristic slowing of resting-state rhythms (elevated theta and reduced alpha and increased gamma). For completeness, we compute TAR as TAR = *P̄_θ_ /P̄_α_*, where *P̄_·_* denotes the mean power within each canonical band. While some works use absolute power, some other relative power or both [85], here, except for TAR we considered the absolute average power for the rest of the frequency bands.
- Sample Entropy (SampEn) for each frequency band, using embedding dimension *m* = 2 and tolerance *r* = 0.2 *σ*, where *σ* is the standard deviation of the corresponding band-limited signal. SampEn quantifies regularity by estimating the likelihood that two length-*m* sequences that match within the tolerance remain similar when extended to length *m*+1. By excluding self-matches, SampEn reduces bias relative to Approximate Entropy and is particularly suitable for short or moderately sized time series typical of resting-state EEG windows [86]. This measure is computed independently per band to probe band-specific complexity changes associated with neurodegeneration.
- Fractal dimension measures, Katz Fractal Dimension (KFD) and Higuchi Fractal Dimension (HFD), for each band. KFD provides a compact descriptor of waveform complexity by relating the cumulative curve length to the maximal amplitude excursion, thereby capturing the overall jaggedness of the time series over its duration [87]. HFD, in contrast, estimates how the effective curve length scales as the sampling interval is varied, yielding a multiscale assessment of irregularity and self-similarity that is sensitive to fine temporal structure in EEG [88]. Using both indices provides complementary views of scale-dependent complexity. Fractal-dimension measures, have been shown to effectively discriminate between different brain states and pathological conditions in EEG recordings [89, 90, 91].
- The Hurst exponent, *H*, which quantifies the degree of long-range temporal dependence in the time series [92]. Values *H >* 0.5 indicate persistence (positive long-memory correlations), *H <* 0.5 indicates antipersistence, and *H* ≈ 0.5 is consistent with uncorrelated dynamics. Estimating *H* per band enables us to assess whether dementia-related slowing is accompanied by systematic changes in long-range dependence.
- The family of complexity metrics, Extrema-Segmented Entropy (ExSEnt), [35], applied per band and described in detail in the next subsection. In brief, ExSEnt segments the signal using local extrema and computes entropy-based descriptors over extrema-defined segments, providing complementary sensitivity to amplitude- and timing-related irregularities that are not fully captured by conventional entropy or fractal indices.

Apart from the power, the rest of the features were not computed over the delta band, since the slow oscillations of this frequency over a short range of 23 seconds do not provide rich information for irregularity evaluation.

### 2.5. ExSEnt: a novel set of temporal and amplitude-driven entropy metrics

The ExSEnt quantifies signal complexity from both temporal and amplitude perspectives. After segmenting the data based on successive local extrema, this method quantifies the complexity of durations and net amplitudes of these segments. The procedure unfolds in three stages. First, the signal is segmented based on local minima and maxima: each interval between consecutive extrema defines a segment (Fig. 3). From each segment, two features are extracted—the duration and the signed amplitude change:

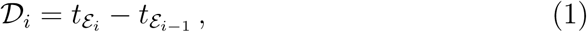

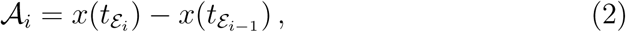

**Figure 3:**
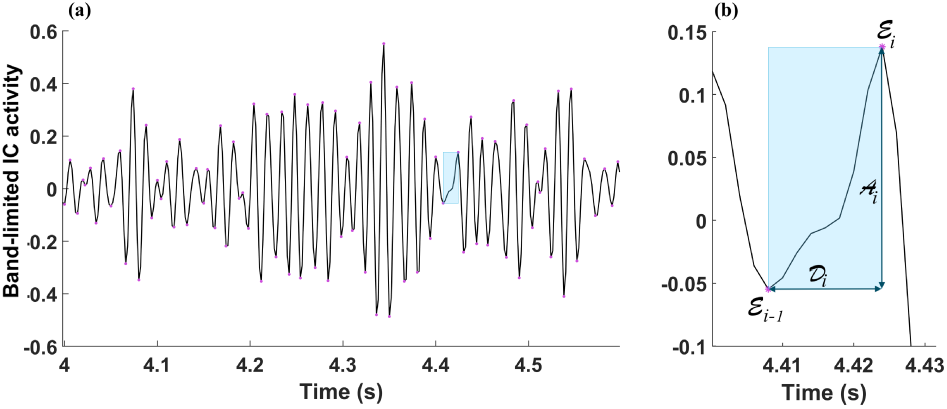
Illustration of the extrema-based segmentation underlying ExSEnt metrics over a low-gamma EEG trace from an AD patient. Panel (a) displays a segment of the signal, with one interval between two consecutive extrema highlighted. Panel (b) shows a zoomedin view of this interval, illustrating how ExSEnt metrics are derived: A*_i_* denotes the net amplitude change across the segment, while D*_i_* represents its duration.

where ℰ*_i_* is the *i^th^* extreme point. After extracting these features for all the segments, SampEn is computed separately for the sequences {𝒟*_k_*} and {𝒜*_k_*}, yielding entropies ℋ*_𝒟_* and ℋ*_𝒜_* that reflect the irregularity in segment durations and amplitudes, respectively:

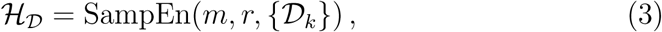

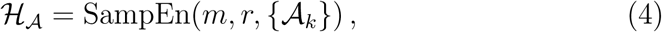

where *m* is the embedding dimension and *r* the tolerance. The tolerance specifies the maximum allowable difference (usually computed as the Chebyshev distance) for two sequences to be considered similar and is typically set to 20% of the signal’s standard deviation.

Finally, SampEn is computed for the paired sequence {(𝒟*_k_,* 𝒜*_k_*)}, resulting in a joint entropy ℋ*_𝒟𝒜_* that captures the degree of coupling between temporal and amplitude variations:

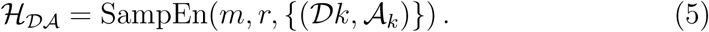

Lower ℋ*_𝒟𝒜_* indicates stronger synchronization between duration and amplitude features, while higher values suggest more independent and complex interactions. This approach yields three complementary entropy metrics, each capturing distinct aspects of local extrema dynamics. Additionally, we quantified the number of extracted segments per window (*M*), which serves as a supplementary complexity indicator by reflecting the signal’s local volatility and segmentation density.

Thus, we extracted nine groups of features across five frequency bands, together with the TAR, yielding a total of 46 features. In addition, we compute the delta-band (1–4 Hz) signal mean, bringing the count to 47 features over 23 windows for each subject. To obtain a single subject-level representation suitable for stability selection, we aggregate each feature across windows using robust summary statistics—median, interquartile range (IQR), and coefficient of variation (CV = SD*/*mean). These summaries mitigate the impact of outliers and inter-trial variability that are common in EEG, especially when comparing across subjects. Concatenating the three summaries for each feature produces 47 × 3 = 141 subject-level features.

High-dimensional feature spaces often contain redundant or noisy predictors that can impair generalization. We applied stability selection to identify the most stable and informative variables [48, 43, 93]. To this end, we implemented a Leave-One-Subject-Out (LOSO) cross-validation scheme, ensuring subject-level independence, and embedded stability selection with penalized LR to extract features that were consistently selected across resampling iterations. The following subsections detail this procedure, including the stability-selection framework, the nested LOSO design, and the aggregation of results across folds.

### 2.6. Stability selection

Stability selection identifies the variables that most frequently contribute to class separation in randomly drawn subsets of the data. It is particularly well suited to a high-dimensional design matrix, *X_n×p_*, where the number of features, *p*, exceeds the number of samples, *n*; in the case of the current dataset, the number of samples in the selected brain areas is *n*_LPFC_ = 45, *n*_LVA_ = 55, *n*_RPFC_ = 58, and *n*_RVA_ = 67, and the number of features is *p* = 141.

Hence, by performing stability selection, we aim to retain the features that are most stable in separating the two classes (healthy vs. dementia). We used LR with elastic-net penalty, parameterized by the *ℓ*_1_ mixing ratio (*ℓ*_1_ ∈ [0, 1]):

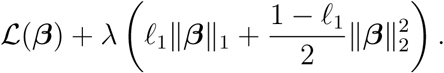

In our Python implementation, we used scikit-learn’s LogisticRegression, where the regularization parameter is expressed as *C* ∝ 1*/λ*, so we will refer to the regularization coefficient by *C*. Thus, a larger *C* corresponds to weaker regularization and typically yields less sparse (denser) solutions and vice versa. Elastic net is well-suited for stability selection since it balances sparsity with grouping effects that help when features are correlated [94, 43].

Given *B* subsamples of size *τn* (*τ* ∈ (0, 1) fraction of total samples) and a grid of regularization strengths {*λ*}, we fit the model on each subsample and recorded a binary indicator *I_jb_*denoting whether feature *j* is selected in run *b*. Therefore, the selection frequency for feature *j* is

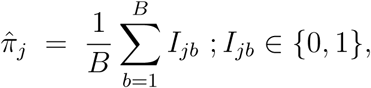

and the stable set is defined as S = {*j* : *π̂_j_* ≥ *π*_thr_ }, where *π*_thr_ is the selection-frequency threshold.

We performed classical stability selection with half-sampling (*τ* = 0.5) to adhere to the Meinshausen–Bühlmann (MB) framework, for which control of the per-family error rate (PFER) is available in closed form. PFER is defined as the expected number of falsely selected features among the retained set. Under mild conditions, the PFER is bounded by

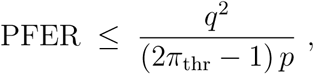

where *p* denotes the total number of candidate features, *q* is the average number of selected features per subsample [95]. The selection threshold *π*_thr_ (along with PFER and the elastic net mixing parameter) was determined systematically using the grid-search procedure described in subsection 2.9.

To address class imbalance, we applied inverse class-frequency weighting, giving proportionally higher weight to the minority class so that both groups contributed equally to the loss function during model training. The optimization was allowed to run for up to 5000 iterations with a convergence tolerance of 10*^−^*^4^, meaning that the optimization terminated once the improvements in the objective function fell below this threshold. This relatively strict criterion helps prevent premature stopping. It ensures that the solution has adequately converged, which is particularly important when using strong regularization, where coefficients can be small and require finer optimization steps to stabilize. Nonzero coefficients defined the selected features (*>* 10*^−^*^6^). The regularization strength *C* was tuned via a pilot grid search on logarithmically spaced values to achieve a sparse model that satisfied the target false selection bound, PFER, at the stability threshold, *π*_thr_. In practice, this ensured that each subsample retained only a small fraction of predictors with nonzero coefficients, promoting sparsity while maintaining coverage of informative features. By tuning *C* to meet the target sparsity implied by the PFER bound, the procedure favored a compact yet informative subset of features, enhancing both interpretability and stability.

### 2.7. Nested model selection and calibrated decision rule

In order to perform stability selection, the model must be repeatedly trained on subsampled versions of the training set so that features consistently contributing to classification can be distinguished from those selected only by chance. To achieve this, we implemented a nested model selection in each LOSO fold. To do so, the LR was fitted twice, for two different purposes:

1. *Nested stability selection*: we fit an elastic net to perform stability selection and extract the features from training data.
2. *Classification*: we used selected features and fitted a model with *ℓ*_2_ regression on the training set, which was used to predict the probability of dementia for the left-out subject of the LOSO fold.

We used elastic net only for feature selection because it yields a sparse, correlation-aware set of predictors under subsampling, and then refit an *ℓ*_2_-regularized logistic model on the selected features because ridge regularization shrinks coefficients smoothly (without setting them to zero), which reduces variance and sensitivity to small data perturbations when predictors are highly correlated, typically improving out-of-sample generalization and producing more stable coefficient estimates.

In the stability selection step, we generated stratified subsamples of the data, drawn without replacement, to preserve class balance while introducing variability across iterations. Each subsample was then normalized via z-score or robust normalization to ensure comparable scales across variables. Normalization was performed with normalization parameters estimated from the training set only and then used for normalizing both the training and test sets. Logistic regression with elastic net was subsequently fitted on each subsample, and the set of features with non-zero coefficients was recorded. Repeating this procedure across many subsamples yields a stable estimate of how frequently each feature is selected. A presentation of the end-to-end processing steps along with the detailed nested-stability selection are presented in Figs. 4 and 5.

**Figure 4:**
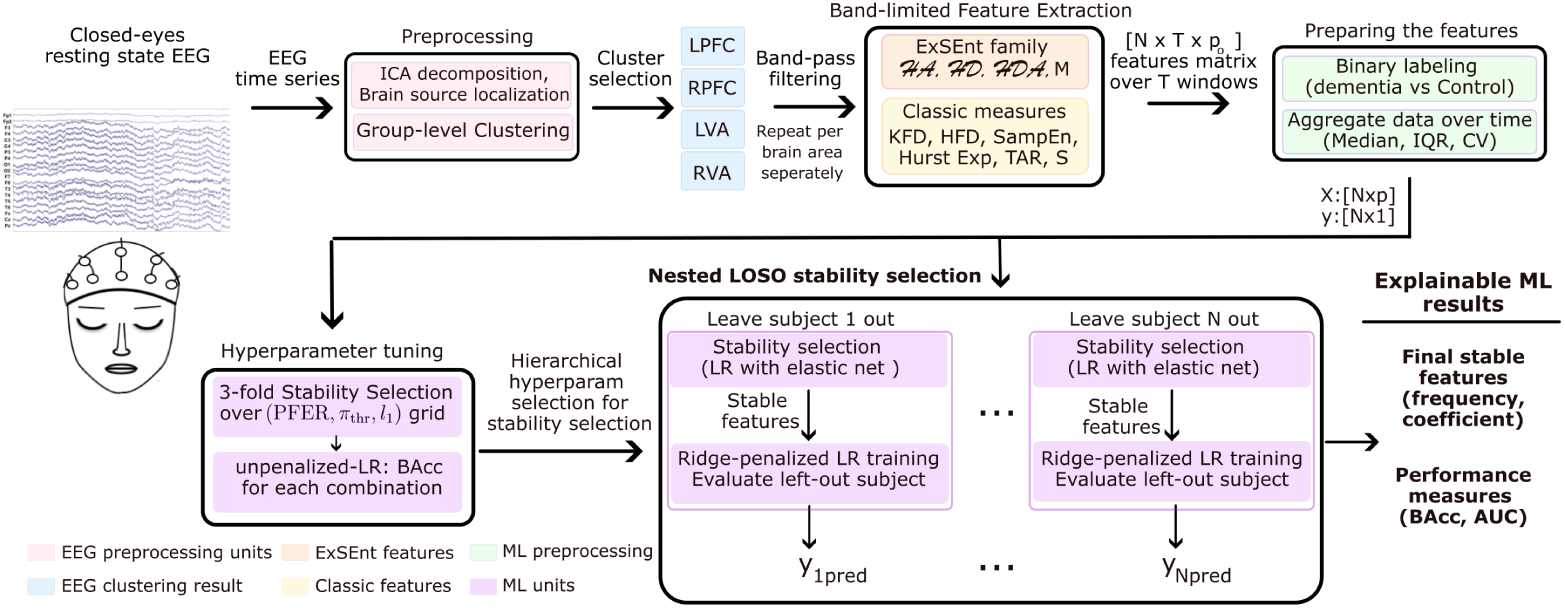
End-to-end overview of the analysis pipeline from EEG recording to classification. After preprocessing, ICA-based source decomposition, source localization, and group-level clustering, four brain-area clusters were selected and processed separately. For each area, band-pass filtering was applied and band-limited features were extracted over 23 time windows, forming an *N* × *T* × *p*_0_ feature tensor. Features were then aggregated across windows using summary statistics and paired with binary labels (dementia vs. control) to yield (**X** ∈ ℝ*^N×p^,* **y** ∈ ℝ*^N×^*^1^). Stability-selection hyperparameters (PFER*, π*_thr_*, ℓ*_1_) were chosen by evaluating all grid combinations via 3-fold stability selection, and were subsequently used in a nested LOSO scheme: within each fold, elastic net stability selection identified stable features, an *ℓ*_2_-penalized logistic model was trained over them, and the left-out subject was evaluated subsequently. The output comprised final stable features (selection frequency and coefficients) and performance metrics (BAcc, AUC).

**Figure 5:**
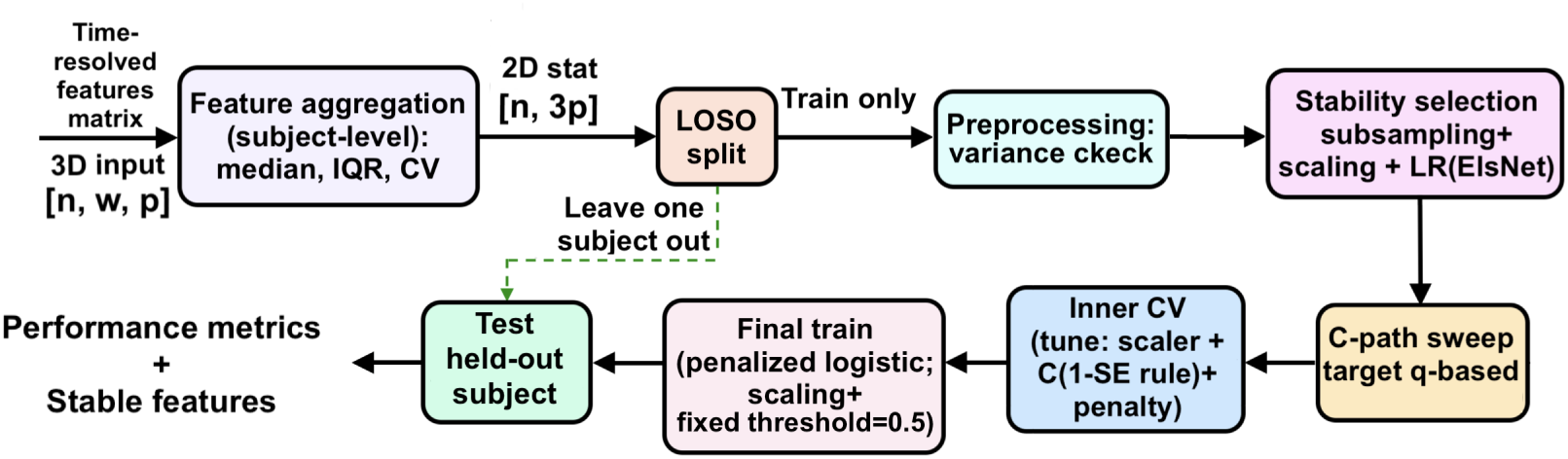
Block diagram of the stability-selection and nested model-selection procedure. The 3D input matrix contains time-resolved feature values, where *n*, *w*, and *p* denote the number of subjects, sliding windows, and features, respectively. After aggregating features at the subject level (using median, IQR, and CV), a LOSO scheme was applied. For each training fold, preprocessing, stability selection, and inner cross-validation were performed to identify the optimal regularization strength *C* and scaling strategy based on mean balanced accuracy (BAcc), using the 1-SE rule. A fixed decision threshold of 0.5 was used for classification. The range of *C* values was adaptively adjusted for each brain area to ensure stable convergence and avoid over- or under-regularization. Final metrics and stable features were derived from LOSO evaluations.

To perform stability selection, we used a predefined set of hyperparameters (selection frequency threshold *π*_thr_, PFER, *l*_1_ ratio, and *C* grid, as described in subsection 2.9). In each stability-selection run, after drawing *B* stratified subsamples of size ⌊*τn*⌋ per class, we fitted an elastic net over the grid of regularization strengths *C*, and adaptively chose the *C* value that yielded a number of nonzero coefficients which was closest to the *q*_target_, according to MB:

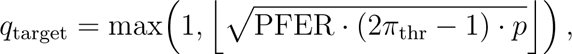

so that the corresponding stability-selection PFER upper bound could be controlled in expectation. The resulting per-feature selection frequencies were computed across subsamples, and the candidate set was defined as all features with frequency ≥ *π*_thr_. If no feature exceeded *π*_thr_ in a given run, we enforced a non-empty candidate set by retaining the top-*m* features ranked by frequency, with *m* set to the mean number of selected features across subsamples (*q*^).

After this, in the classification step, each candidate feature set was evaluated in an inner 3-fold cross-validation on the training data. We compared two scaling strategies as before and swept a logarithmic grid of *C* values for the final *ℓ*_2_-penalized LR. *C* was selected using the one–standard–error (1-SE) rule: we chose the smallest *C* whose mean balanced accuracy (BAcc) was within one standard error of the best mean BAcc, thereby favoring stronger regularization. Finally, the complete pipeline was refit on the full training set using the selected features and applied once to the held-out subject.

### 2.8. Evaluation and across-fold stability

The performance was computed from the LOSO held-out predictions, reporting BAcc for the default classification threshold *θ*_thr_ = 0.5, and for the post-hoc and the nested LOSO Youden thresholds, as well as the Area Under the ROC Curve (AUC) obtained from the pooled held-out probabilities. The primary operating point throughout model selection and reporting was the fixed threshold of 0.5, ensuring a consistent decision rule across folds and avoiding threshold optimization within the training loop. For completeness, we computed two Youden BAcc thresholds. First, a nested Youden threshold was estimated independently within each outer LOSO fold using only the training subjects (e.g., via an inner cross-validation) and then applied to the held-out subject, thereby preventing test-label leakage during threshold selection. Second, we computed a pooled post-hoc Youden threshold on the pooled LOSO test probabilities; because this threshold is tuned directly on the evaluation outputs (pooled test-fold probabilities and labels), it is optimistically biased, was not used for model selection, and is reported only as an optimistic upper bound.

Feature robustness was quantified using two complementary notions of stability. First, we tracked the across-fold retention rate of each feature in the final LOSO models (outer selection frequency). Second, for each fold we retained the full stability-selection frequency vector of the chosen stability-selection run and summarized it across folds by its mean and minimum frequency, providing a direct measure of selection consistency under repeated subsampling. A global stable set was defined by thresholding the mean stability-selection frequency across folds, capped to the top-20 features, yielding a compact subset with high subsampling-level reproducibility.

### 2.9. Hyperparameter tuning for stability selection sparsity control

Before performing the nested LOSO as explained above, we had to select the hyperparameter values. This was specifically important because in the stability selection, the values of PFER and *π_thr_* have to be fixed for all the folds, so it is not possible to include a range of values and do a grid search while running the stability selection for biomarker evaluation. To select the sparsity-control parameters, we performed an explicit grid search. For each brain area, we evaluated all combinations of triplet (PFER*, π*_thr_*, l*_1_) in a threedimensional grid:

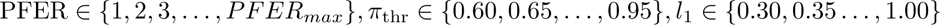

Here, PFER_max_= ⌈0.1 *p*⌉, thus PFER_max_*_|_*_WE_ = 15 and PFER_max_*_|_*_WoE_ = 9. Each triplet (PFER*, π*_thr_*, l*_1_) was evaluated with stratified 3-fold cross validation. Stability selection was performed with *B* = 500 stratified subsamples at subsample fraction *τ* = 0.5. Regularization strengths were 30 values, logarithmically spaced over 10*^−^*^5^ to 10^3^, in 3-fold stability selection (hyperparameter tuning step Fig. 4).

Then, we fitted an unpenalized LR model to the stability-selected features and computed the balanced accuracy (BAcc). Here as well, both standard and robust scaling were evaluated. For each fold, the best of the two scalers based on BAcc was retained, and the fold-wise best BAcc values were averaged across folds. We also logged the set of *C* values selected across subsamples, and the number of retained features per fold.

After completing the hyperparameter grid search, we applied an explicit selection policy to choose a single configuration for subsequent LOSO analysis. For each triplet, we summarized cross-validation outcomes across folds. A fold was considered valid if it had retained at least one feature (*q >* 0). For each triplet we computed the mean BAcc across valid folds, its standard error (SE), the number of valid folds, the median number of selected features, and the IQR of the selected feature count. In addition, we reported the effective regularization range used by the stability-selection base learner by extracting the minimum and maximum of the unique *C* values selected across subsamples (*C*_min_ and *C*_max_), obtained from the stored grid-search outputs. Among eligible configurations, we identified the maximum BAcc and defined a near-optimal band, 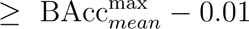 to avoid over-interpreting negligible cross-validation differences.

Within these configurations, we promoted interpretability and stability by applying a compactness window on the median feature count (1 ≤ *q̂_med_* ≤ ⌈0.1 *p*⌉) and a cross-fold variability constraint (*q̂*_IQR_ ≤ 2), relaxing these only if infeasible. Final selection used a hierarchical ordering of candidate hyperparameter sets, ranking by: *(i)* higher BAcc*_mean_*, ii. lower *q*_IQR_, iii. Lower *q̂*_med_, iv. lower PFER, v. higher *π*_thr_, with *l*_1_ used only as a final tie-breaker (minimum distance to 0.6). As a ranking-sensitivity check, we repeated the selection after swapping the priority of *π*_thr_ and PFER.

For each top-ranked hyperparameter setting, we performed the stability selection with LOSO, using the top triplet and the *C* range based on extracted *C*_min_ and *C*_max_ values. We set the range of *ℓ*_2_ to a wide range. The aim of this LOSO was to extract the stability-selected features of each fold. We used these per-fold selected and pre-saved features to perform a final pooled LOSO fit of the *ℓ*_2_-regularized classifier over the candidate *C* grid. We then selected a stable regularization level using a Δ-plateau criterion on pooled LOSO balanced accuracy (Δ = 0.01): all *C* values within Δ of the maximum BAcc were considered acceptable, and we chose the most conservative representative value *C*_rep_. All reported metrics and biomarker specifications were computed at this fixed *C*_rep_.

## 3. Results

To assess our main hypothesis—the contribution of the proposed ExSEnt metrics to the classification performance—we performed the full stability-selection procedure described in Section 2.6, once on the full set (WE) and once excluding the ExSEnt family measures (WoE). Classification was conducted independently for each of the four brain areas (LVA, RVA, LPFC, RPFC) under both feature sets. This design enables a direct, within-cluster comparison that quantifies the additive value of ExSEnt relative to the remaining nonlinear and spectral feature families. In the following section, we first report the hyperparameter-tuning outcomes used to select a single stability-selection configuration per cluster and feature set, and then present the corresponding nested LOSO results.

### 3.1. Hyperparameter tuning results

We established the stability-selection hyperparameters via a grid search over (PFER*, π*_thr_*, l*_1_) using 3-fold cross-validation and 500 subsamples. This tuning stage was used to identify configurations that were well-defined across folds (no empty selections) and that jointly balanced predictive performance and biomarker compactness. For each candidate triplet, we summarized mean BAcc and sparsity statistics (median number of selected features and IQR) across folds, and retained the top-ranked configurations for each cluster separately for the feature sets with and without ExSEnt. These tuning-stage summaries provide an initial comparison of how ExSEnt affects performance and sparsity under identical stability-selection constraints, and they define the hyperparameter settings subsequently used in the nested LOSO evaluation.

Across clusters, the effect of including ExSEnt during the initial tuning stage was homogeneous (Table 2, shown for the hierarchical policy prioritizing PFER before *π*_thr_). All the clusters showed higher performance when ExSEnt was included. In the LPFC, the configurations including ExSEnt achieved markedly higher BAcc than configurations excluding ExSEnt, with comparable sparsity. In RPFC, we had the next highest increase by including ExSEnt. In LVA, including ExSEnt produced a slightly higher mean BAcc than excluding it. In RVA, including ExSEnt improved mean BAcc, with a median feature equal to 1, while excluding it led to a larger median number of selected features.

**Table 2:** Stability-selection hyperparameters and cross-validation summaries with and without ExSEnt, top 3 sets per cluster; hierarchical policy prioritizing PFER before *π*_thr_. Reported *C* range is [*C*_min_*, C*_max_] extracted from the stored stability-selection runs.

| Cluster | With ExSEnt |  |  |  | Without ExSEnt |  |  |  |
| --- | --- | --- | --- | --- | --- | --- | --- | --- |
| | (PFER, $\pi_{thr}$, $l_1$) | BAcc | $\hat{q}[\hat{q}_{IQR}]$ | $C$ range | (PFER, $\pi_{thr}$, $l_1$) | BAcc | $\hat{q}[\hat{q}_{IQR}]$ | $C$ range |
| LPFC | (10,0.65,0.30) | **0.841** | 2 [2] | [0.073,0.924] | (8,0.70,0.75) | 0.439 | 1 [0.5] | [0.489,22.122] |
|  | (11,0.60,0.30) | 0.841 | 2 [2] | [0.073,0.924] |  |  |  |  |
| RPFC | (5,0.60,0.35) | **0.664** | 1 [0.5] | [0.073,0.259] | (8,0.75,0.65) | 0.581 | 2 [0.5] | [0.489,11.721] |
| LVA | (5,0.70,0.30) | **0.780** | 3 [3] | [0.073,0.259] | (5,0.75,0.65) | 0.779 | 2 [0.0] | [0.259,6.210] |
|  | (6,0.60,0.35) | 0.780 | 3 [2.5] | [0.073,0.259] | (6,0.80,0.60) | 0.779 | 2 [0.5] | [0.259,6.210] |
|  | (7,0.65,0.30) | 0.780 | 3 [2.5] | [0.073,0.259] | (7,0.75,0.95) | 0.780 | 2 [0.5] | [0.924,41.753] |
| RVA | (13,0.80,0.30) | **0.626** | 1 [1] | [0.137,1.743] | (8,0.70,0.30) | 0.592 | 3 [1.5] | [0.073,0.259] |

In all cases, the selected sets were typically very small despite comparatively larger tuned PFER values. This was not unexpected, because PFER was an upper-bound–type control parameter tied to the average sparsity of the base elastic net selector across subsamples, whereas *q*^ reported only the features that ultimately exceeded the stability threshold. In our setting, redundancy across summary statistics and frequency bands induced correlated feature groups, so selection probability was often distributed across near-substitutes: many features may have been chosen intermittently in subsamples but failed to reach *π*_thr_, leaving only a few representative predictors in the final set. In addition, the PFER bound was known to be conservative (and can be loose under correlated predictors), so it can overstate the expected number of false discoveries relative to the realized stable set size.

We repeated the selection after swapping the priority of *π*_thr_ and PFER. With this swap, the selected configuration remained unchanged for most clusters; differences were observed only in LVA, both with and without ExSEnt. The corresponding top-ranked configurations under the swapped ordering are summarized in Table 3, for direct comparison with the primary ordering. In the rest of the study, we focused on the top configurations from Table 2, since the difference between the two settings was not significant.

**Table 3:** Stability-selection hyperparameters and cross-validation summaries, the same as Table 2: by hierarchical policy prioritizing *π*_thr_ before PFER.

| Cluster | With ExSEnt |  |  |  | Without ExSEnt |  |  |  |
| --- | --- | --- | --- | --- | --- | --- | --- | --- |
| | (PFER, $\pi_{thr}$, $l_1$) | BAcc | $\hat{q}[\hat{q}_{IQR}]$ | $C$ range | (PFER, $\pi_{thr}$, $l_1$) | BAcc | $\hat{q}[\hat{q}_{IQR}]$ | $C$ range |
| LVA | (5, 0.70, 0.30) | **0.780** | 3 [3] | [0.073, 0.259] | (6, 0.80, 0.60) | 0.779 | 2 [0.5] | [0.259, 6.210] |
|  | (7, 0.65, 0.30) | 0.780 | 3 [2.5] | [0.073, 0.259] | (7, 0.75, 0.95) | 0.780 | 2 [0.5] | [0.924, 41.753] |
|  | (6, 0.60, 0.35) | 0.780 | 3 [2.5] | [0.073, 0.259] | (8, 0.70, 1.00) | 0.780 | 2 [0.5] | [0.924, 41.753] |

### 3.2. Nested LOSO results

Using the tuning-stage ranking in Table 2, we fixed the top stability-selection configuration (PFER*, π*_thr_*, l*_1_) for each brain area and feature set (WE/WoE), and evaluated it in a nested LOSO protocol. In each outer LOSO fold, stability selection was estimated using only the training subjects, with 1000 subsampling—from the training set, we subsampled half of the data, for 1000 without replacement, while making sure we have at least 2 subjects from each class—, ensuring that feature selection remained strictly out-of-sample with respect to the held-out subject. The inner loop served to produce well-defined feature sets across folds, which met the *π_thr_* and remained close to the target sparsity, *ℓ*_1_.

A practical consideration was specifying the inverse regularization parameter *C* for LR, which we found to have a non-negligible impact on predictive performance. In our pipeline, *C* intervenes at two distinct stages. First, during nested stability selection, *C* controls the regularization strength of the elastic net base estimator; sweeping *C* over the range yields the regularization (sparsity) path used within subsampling. For each brain-area/feature-set configuration, we reused the performance-consistent bounds [*C*_min_*, C*_max_] obtained in the tuning stage (Table 2). To construct the stability path employed in the nested LOSO runs, we used these bounds in log_10_(*C*) and discretized the resulting interval at a fixed resolution of 20 points per decade (bounded to [20, 500] grid points), yielding *C*_STAB_.

Second intervention of *C* was in the final evaluation with the *ℓ*_2_ regularization, where we fitted the model to the fold-selected features. To quantify the sensitivity of the final refit to *C* (independently of stability selection), we performed a pooled LOSO evaluation over a global grid *C*_FINAL_ = logspace(−5, 4*, n*) with the same density (20 points) per decade. Importantly, for each outer fold, we fixed the feature subset selected from the training subjects only during stability selection, and then refit an *ℓ*_2_-regularized logistic-regression model on the corresponding training set for each *C* ∈ *C*_FINAL_ to obtain the predicted class probability of the held-out subject. Pooling these left-out probabilities across folds yielded BAcc curves as a function of log_10_(*C*). The results depicted in Fig. 6 showed that overall performance exhibited broad plateaus as a function of log_10_(*C*) in many of the intervals, indicating that the final refit is not critically dependent on a single *C* value; nevertheless, the effect was non-negligible in several configurations.

**Figure 6:**
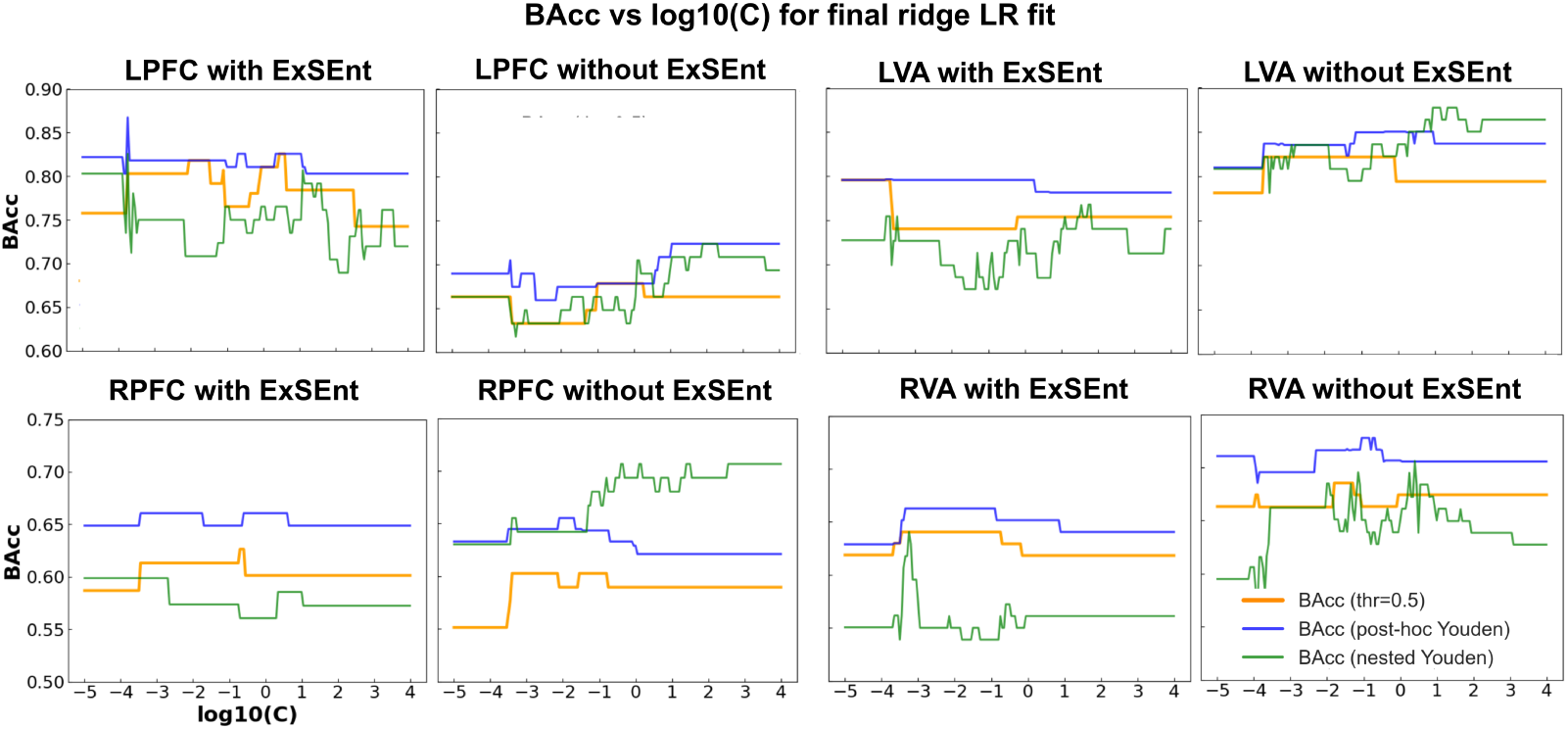
Balanced accuracy as a function of the inverse regularization parameter, log_10_(*C*), for *ℓ*_2_-regularized LR across four brain regions (LPFC, RPFC, LVA, RVA). For each region, results are shown using the full feature set, including ExSEnt (with ExSEnt) and after removing ExSEnt (without ExSEnt). Three decision-threshold strategies are compared: fixed threshold (0.5), post-hoc Youden threshold, and nested Youden threshold (estimated within the cross-validation loop).

For the LPFC with ExSEnt, BAcc at threshold 0.5 ranged approximately from 0.74 to 0.83, whereas the corresponding LPFC model without ExSEnt showed a lower performance. The LVA curves were comparatively flat, suggesting higher robustness of the final refit to regularization in that region. Right-hemisphere clusters displayed lower absolute performance and larger variability with *C*. The RPFC area showed a similar range of performance in both sets, while RVA performed slightly better across the *C* without ExSEnt. Across all panels, post-hoc Youden thresholding mostly yielded higher BAcc than the fixed threshold (optimistic by construction), while the nested Youden curves showed increased fluctuations due to per-fold threshold estimation on training data only.

Following the pooled-LOSO sensitivity analysis (Fig. 6), we extracted for each configuration a near-optimal regularization interval from the BAcc(*C*) curve by selecting the Δ-plateau, defined as the set of *C* values satisfying BAcc(*C*) ≥ max*_C_′* (BAcc(*C^′^*)) − Δ with Δ = 0.01, and denoted its bounds by [*C*_lo_*, C*_hi_]. We then summarized the corresponding pooled-LOSO discrimination performance within this plateau in Table 4. The corresponding values for AUC and BAcc at threshold 0.5, together with Youden-based balanced accuracy computed either post-hoc on the pooled LOSO scores (YBAcc*_post_*) or via nested, training-only threshold estimation (YBAcc), are presented in Table 4 thereby linking the plateau-based *C* selection to the final reported performance while avoiding reliance on a single finely tuned *C* value.

**Table 4:** AUC, Balanced accuracy (Acc), nested Youden Balanced accuracy (YBAcc), and post-hoc Youden Balanced accuracy (YBAcc*_post_*) for the four brain areas with and without inclusion of ExSEnt metrics.

| Cluster | With ExSEnt |  |  |  | Without ExSEnt |  |  |  |
| --- | --- | --- | --- | --- | --- | --- | --- | --- |
| | AUC | BAcc | YBAcc | $\text{YBAcc}_{\text{post}}$ | AUC | BAcc | YBAcc | $\text{YBAcc}_{\text{post}}$ |
| LPFC | 0.854 | **82.6%** | 82.6% | 86.7% | 0.678 | 71.5% | 72.3% | 72.3% |
| LVA | 0.832 | 79.6% | 76.8% | 79.7% | 0.880 | 82.2% | 85.0% | 87.7% |
| RPFC | 0.692 | 62.6% | 59.8% | 66.1% | 0.649 | 60.2% | 70.7% | 65.5% |
| RVA | 0.610 | 64.0% | 64.0% | 66.3% | 0.771 | 68.5% | 70.6% | 72.8% |

Figure 7 reports pooled LOSO receiver-operating-characteristic (ROC) curves for each brain-area cluster, shown for both WE and WoE. For each panel, the ROC was computed from the pooled out-of-fold probabilities obtained at a fixed regularization value *C* selected from the evaluation grid (the value shown in the panel header), and the corresponding AUC is reported together with a 95% confidence interval estimated via a subject-level, class-stratified bootstrap (resampling subjects with replacement within each class). The solid curve depicts the bootstrap median ROC, while the shaded band shows the IQR of the bootstrapped true-positive rate at each false-positive rate, providing an uncertainty envelope around the empirical operating characteristics. The diagonal dashed line indicates chance-level discrimination. In addition, the orange marker denotes the operating point defined by the pooled-LOSO Youden threshold computed from the same left-out probabilities at the chosen *C*.

**Figure 7:**
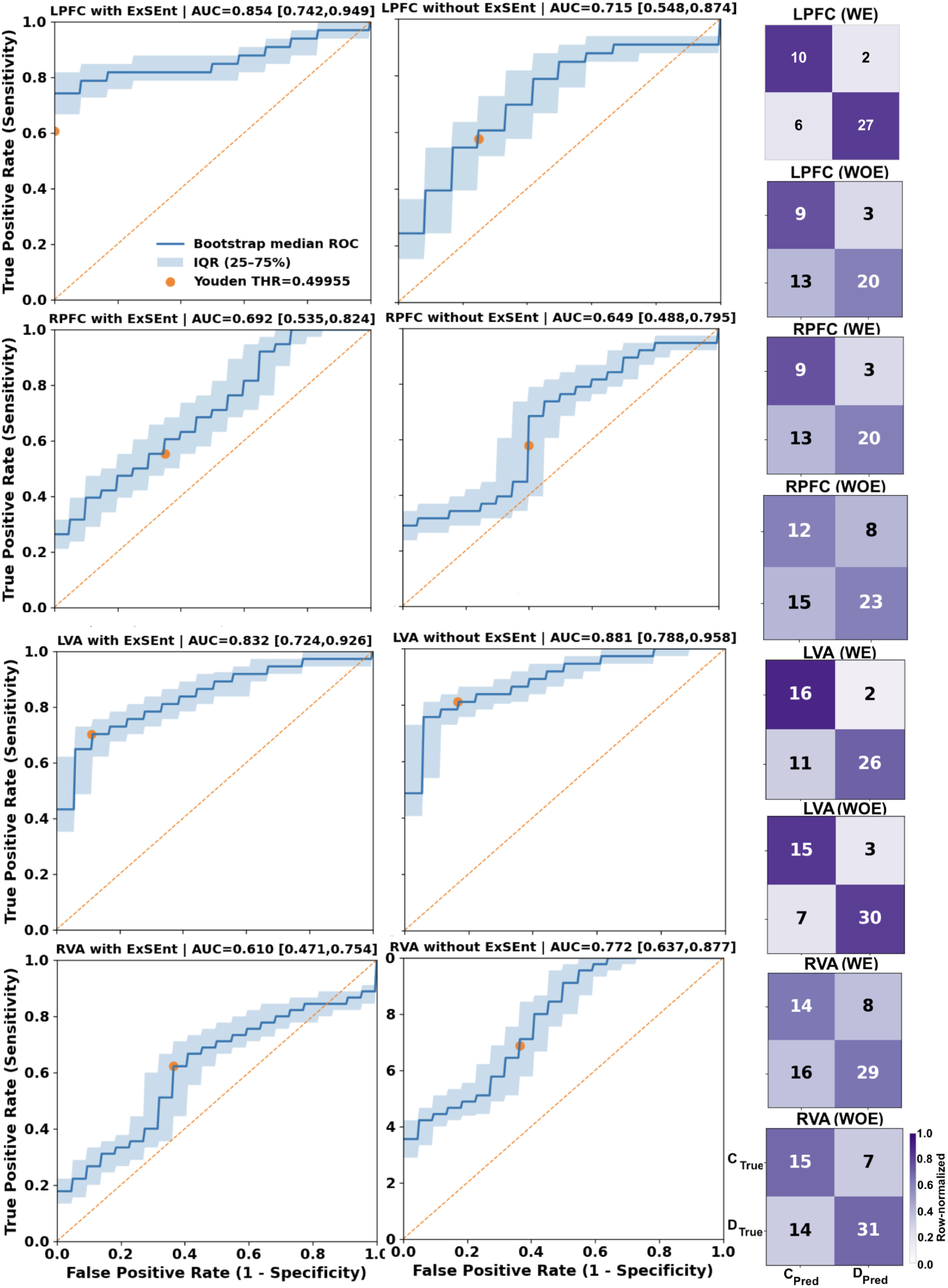
Left: Classification performance; pooled LOSO ROC curves for WE and WoE. Solid curves denote the bootstrap median ROC, and the shaded bands show the interquartile range; the dashed diagonal indicates chance level. Each ROC panel reports the selected regularization strength *C* and the AUC with its 95% bootstrap confidence interval; the orange marker indicates the operating point at the post-hoc Youden threshold. Right: corresponding confusion matrices (counts) with row-normalized shading (colorbar); C and D denote control and dementia, respectively.

Using this threshold= 0.5, we further visualize classification outcomes via row-normalized confusion matrices (right column), where each cell shows the absolute counts and the color intensity represents the within-row proportion, thereby emphasizing sensitivity (positive-class row) and specificity (negative-class row) independently of class prevalence. Together, the ROC panels and the corresponding confusion matrices provide complementary views of discriminability (AUC), uncertainty (bootstrap variability), and the concrete error profile, enabling direct comparison of WE versus WoE within each brain region.

Figures 8 and 9 summarize, for each cortical cluster, the most stable predictors under the LOSO stability-selection pipeline, shown separately for the WE and WoE. For each condition and region, the upper bar plot ranks the top features by their fold-wise selection frequency during stability selection on the training subjects only, while the lower plot highlights the subset that most consistently attains the largest coefficient magnitudes in the final *ℓ*_2_-regularized logistic-regression refit. Bar color encodes the absolute coefficient magnitude, and diagonal hatching denotes negative coefficients (i.e., features inversely associated with the dementia class, such that higher values correspond to lower predicted dementia probability).

**Figure 8:**
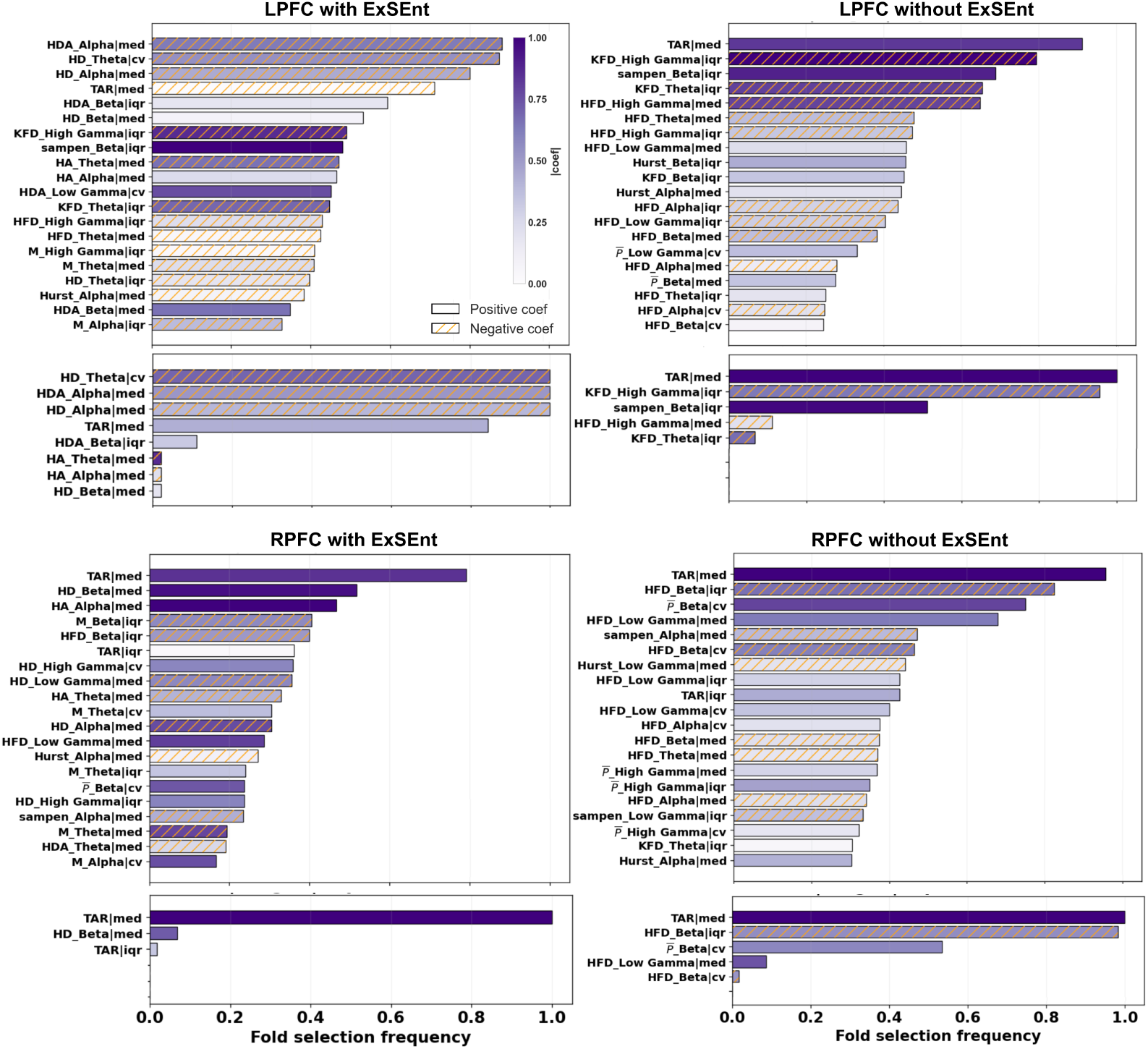
Top 20 stable features selected by the LR model for the right and left PFC areas; with all the features, and excluding the ExSEnt family. For each feature set and brain area, the top plot shows the top features during the stability selection for the training set, and the bottom plot shows the top features in the final LR classification. Bars show the fold selection frequency across leave-one-subject-out runs, color-coded by the absolute coefficient magnitude. Diagonal hatching indicates negative coefficients, while filled bars indicate positive coefficients. Feature labels denote the feature, its frequency band, and the summary statistic used.

**Figure 9:**
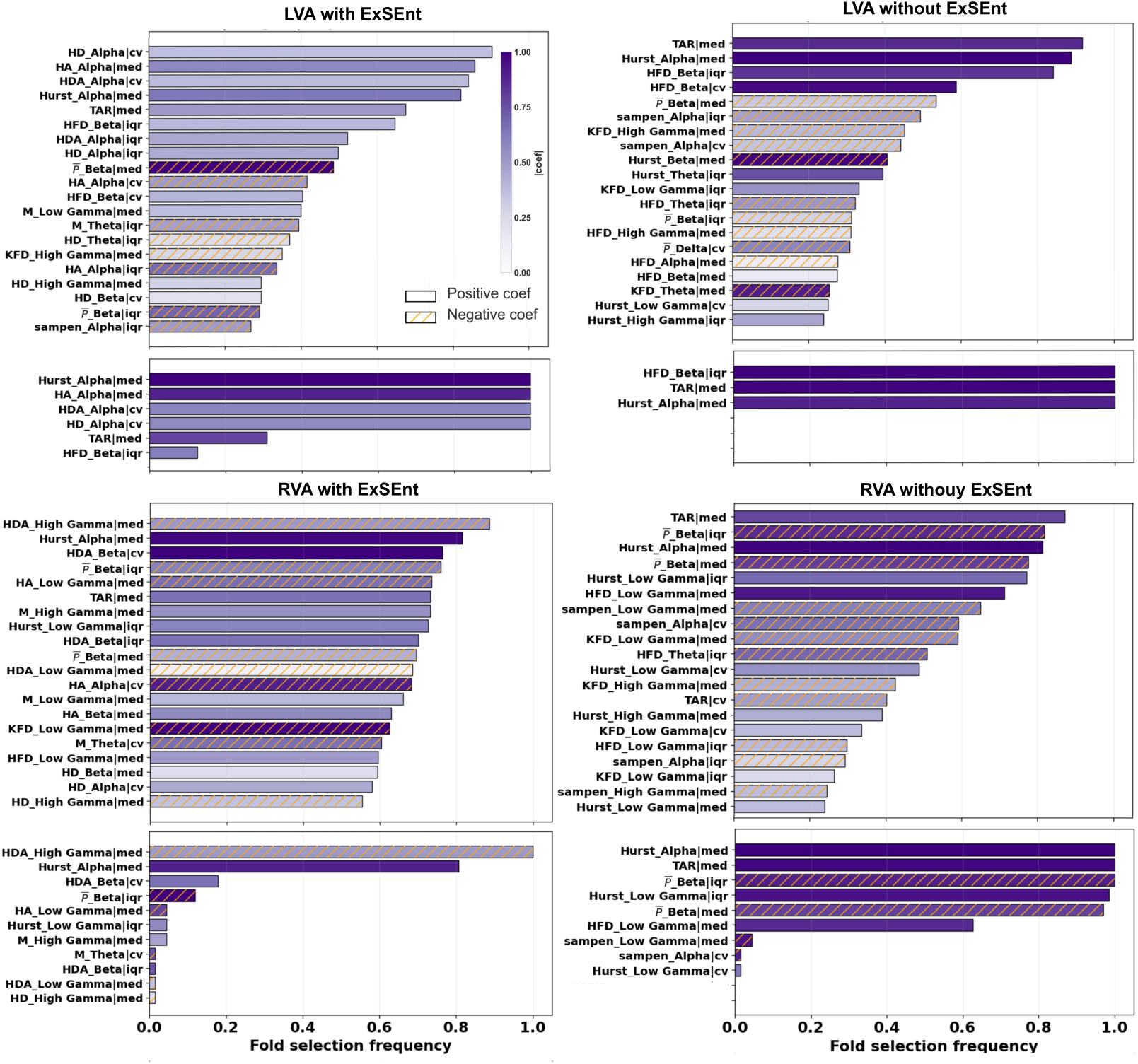
Top 20 stable features selected by the LR model for the right and left VA areas; as in Fig. 8.

We should note that, to enable a consistent coefficient-based ranking across folds, the *ℓ*_2_ refit used for the lower panels is performed on a fixed set of top 20 features obtained from the aggregated stability features. As a result, 20 features are shown in the coefficient plots, ordered by aggregated frequency and absolute value of the coefficient of each feature, whereas Table 2 reports the typical per-fold sparsity implied by the chosen stability-selection hyperparameters. Consequently, while it was expected that only a small subset (e.g., ∼ 2 features per fold) consistently exceeded the *π*_thr_ criterion, a larger set was still refitted under *ℓ*_2_ to yield stable, comparable coefficient magnitudes for visualization.

In the PFC clusters (Fig. 8), LPFC with ExSEnt was characterized by a highly consistent ExSEnt-driven signature: median H*_DAα_*, CV of H*_Dθ_*, and median H*_Dα_* appear with near-unity selection frequency and dominate the final refit, while additional ExSEnt terms across *β* and low-*γ* bands recurred with smaller weights. When ExSEnt metrics were excluded, the LPFC profile shifted toward conventional power, fractal and entropy descriptors, with median TAR emerging as the most stable feature and the final refit relied primarily on it together with KFD (notably high-*γ* IQR) and SampEn (e.g., *β*-band IQR). In RPFC, both conditions were dominated by median TAR, which remained the most stable features and persisted into the final refit; with ExSEnt included, additional ExSEnt measures (e.g., median H*_Dβ_* and H*_Aα_*) entered the stable set but did not replace the TAR/*S̄* as the primary determinants.

In the VA clusters (Fig. 9), LVA with ExSEnt showed a compact, predominantly *α*-band signature in the final refit, driven by ExSEnt features, together with median Hurst*_α_*, while median TAR appeared with lower persistence. After removing ExSEnt, the LVA representation compressed to a small set dominated by median TAR, IQR of HFD*_β_*, and median Hurst*_α_*, which also constitute the final refit. In RVA, inclusion of ExSEnt introduced a high-*γ* ExSEnt component (H*_DA_*) that was highly stable and retained in the final refit followed by median TAR and Hurst exponent median and IQR at *α* and low *γ*, respectively; without ExSEnt, the RVA profile was instead dominated by median TAR and Hurst and *P̄_β_* -, with HFD and SampEn features contributing more intermittently. Overall, these feature-stability profiles provided a mechanistic complement to the performance comparisons, highlighting how inclusion of ExSEnt preferentially reshaped the left-hemisphere models (LPFC and LVA) toward robust ExSEnt-based predictors, whereas right-hemisphere models remained largely driven by TAR and scale-related complexity descriptors.

Table 5 complements Figs. 8 and 9 by providing numerical summaries of feature stability and their coefficients for the most recurrent predictors in each cortical area. For each reported feature *j*, we listed the outer-fold inclusion frequency 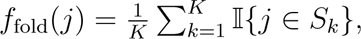 i.e., the proportion of LOSO folds in selection frequency 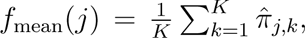 where *π̂_j,k_* was the within-fold subsampling selection probability returned by stability selection; and the signed mean coefficient *w*(*j*) (reported as coef_mean_), computed by averaging the *ℓ*_2_-logistic regression coefficients over those outer folds in which the feature was selected. Thus, *f*_fold_ quantifies cross-fold frequency of the selected feature, *f*_mean_ captures average subsampling-level frequency within training folds, and coef_mean_ summarizes the direction and typical strength of the feature’s coefficient under the fold-specific *ℓ*_2_ refits.

**Table 5:** Top stable features across LOSO folds in all clusters, comparing feature sets with and without ExSEnt. For each feature we report: *f*_fold_, the proportion of LOSO folds in which the feature enters the final fold-specific model, *f*_mean_, the mean stabilityselection frequency across folds, and coef*_mean_*, the average of feature’s coefficients in the fold-specific *ℓ*_2_ refits, averaged over folds in which the feature was selected.

| LPFC (with ExSEnt) |  |  |  | LPFC (without ExSEnt) |  |  |  |
| --- | --- | --- | --- | --- | --- | --- | --- |
| Feature | $f_{\text{fold}}$ | $f_{\text{mean}}$ | $\text{coef}_{\text{mean}}$ | Feature | $f_{\text{fold}}$ | $f_{\text{mean}}$ | $\text{coef}_{\text{mean}}$ |
| $\mathcal{H}_{\mathcal{D}A_{\alpha}}^{\text{md}}$ | 1.000 | 0.880 | -1.723 | $\text{TAR}^{\text{md}}$ | 1.000 | 0.912 | 0.129 |
| $\mathcal{H}_{\mathcal{D}\theta}^{\text{CV}}$ | 1.000 | 0.874 | -1.650 | $\text{KFD}_{\gamma_h}^{\text{IQR}}$ | 0.956 | 0.794 | -0.074 |
| $\mathcal{H}_{\mathcal{D}\alpha}^{\text{md}}$ | 1.000 | 0.801 | -1.061 | | | | |
| $\text{TAR}^{\text{md}}$ | 0.844 | 0.711 | 1.253 | | | | |

| RPFC (with ExSEnt) |  |  |  | RPFC (without ExSEnt) |  |  |  |
| --- | --- | --- | --- | --- | --- | --- | --- |
| Feature | $f_{\text{fold}}$ | $f_{\text{mean}}$ | $\text{coef}_{\text{mean}}$ | Feature | $f_{\text{fold}}$ | $f_{\text{mean}}$ | $\text{coef}_{\text{mean}}$ |
| $\text{TAR}^{\text{md}}$ | 1.000 | 0.792 | $< 10^{-6}$ | $\text{TAR}^{\text{md}}$ | 1.000 | 0.957 | 0.013 |
| | | | | $\text{HFD}_{\beta}^{\text{IQR}}$ | 0.983 | 0.825 | -0.007 |
| | | | | $\overline{P}_{\beta}^{\text{CV}}$ | 0.534 | 0.751 | -0.013 |

| LVA (with ExSEnt) |  |  |  | LVA (without ExSEnt) |  |  |  |
| --- | --- | --- | --- | --- | --- | --- | --- |
| Feature | $f_{\text{fold}}$ | $f_{\text{mean}}$ | $\text{coef}_{\text{mean}}$ | Feature | $f_{\text{fold}}$ | $f_{\text{mean}}$ | $\text{coef}_{\text{mean}}$ |
| $\mathcal{H}_{\mathcal{D}\alpha}^{\text{CV}}$ | 1.000 | 0.903 | 0.043 | $\text{TAR}^{\text{md}}$ | 1.000 | 0.918 | 0.194 |
| $\mathcal{H}_{A_{\alpha}}^{\text{md}}$ | 1.000 | 0.858 | 0.097 | $\text{Hurst}_{\alpha}^{\text{md}}$ | 1.000 | 0.888 | 0.137 |
| $\mathcal{H}_{\mathcal{D}A_{\alpha}}^{\text{CV}}$ | 1.000 | 0.841 | 0.032 | $\text{HFD}_{\beta}^{\text{IQR}}$ | 1.000 | 0.841 | 0.152 |
| $\text{Hurst}_{\alpha}^{\text{md}}$ | 1.000 | 0.820 | 0.076 | | | | |

| RVA (with ExSEnt) |  |  |  | RVA (without ExSEnt) |  |  |  |
| --- | --- | --- | --- | --- | --- | --- | --- |
| Feature | $f_{\text{fold}}$ | $f_{\text{mean}}$ | $\text{coef}_{\text{mean}}$ | Feature | $f_{\text{fold}}$ | $f_{\text{mean}}$ | $\text{coef}_{\text{mean}}$ |
| $\mathcal{H}_{\mathcal{D}A_{\gamma_h}}^{\text{md}}$ | 1.000 | 0.886 | -0.041 | $\text{TAR}^{\text{md}}$ | 1.000 | 0.871 | 0.005 |
| $\text{Hurst}_{\alpha}^{\text{md}}$ | 0.806 | 0.815 | 0.043 | $\overline{P}_{\beta}^{\text{IQR}}$ | 1.000 | 0.818 | -0.004 |
| $\mathcal{H}_{\mathcal{D}A_{\beta}}^{\text{CV}}$ | 0.179 | 0.763 | 0.036 | $\text{Hurst}_{\alpha}^{\text{md}}$ | 1.000 | 0.813 | 0.004 |
| $\overline{P}_{\beta}^{\text{IQR}}$ | 0.119 | 0.759 | -0.256 | $\overline{P}_{\beta}^{\text{md}}$ | 0.985 | 0.776 | -0.004 |

Excluding the ExSEnt family, TAR emerged as the dominant predictor across all regions. In the RPFC, it remained the only selected feature even when ExSEnt measures were available. In contrast, when ExSEnt was included, ExSEnt-derived descriptors constituted the top-ranked predictors-the in rest of the clusters. This pattern was most pronounced in the left-hemisphere, which consistently prioritized ExSEnt measures, whereas the right-hemisphere showed greater reliance on TAR and generally weaker dominance of ExSEnt features.

### 3.3. Subject-level error complementarity

In this subsection, we analyzed the misclassified subjects and quantified cross-model error-set similarity when comparing feature sets within each brain area. Specifically, for each region, we contrasted models trained with ExSEnt features versus without ExSEnt; the corresponding misclassification lists are reported in Tables S1–S8. We then evaluated the overlap between the two error sets using the Jaccard index *J* = |WE ∩ WoE|*/*|WE ∪ WoE|, where a lower *J* indicates greater complementarity (i.e., different subjects were misclassified by the two feature configurations). This subject-wise perspective complements aggregate performance metrics by revealing whether ExSEnt primarily reduced errors on the same difficult individuals (high *J*) or recovered different subjects that were otherwise misclassified (low *J*), thereby clarifying the contribution of ExSEnt to region-specific discriminability.

Table 6 provides a subject-level view of how the two feature sets fail under identical LOSO evaluation. The key observation was that the overlap between misclassified subjects was not complete, implying that WE and WoE representations encoded partially distinct decision evidence. This complementarity was strongest in RPFC, moderate in LPFC and RVA, and limited in LVA, where both models largely misclassified the same subjects and therefore shared similar failure modes.

**Table 6:** Subject-wise complementarity between pooled-LOSO classifiers trained *with* ExSEnt (WE) and *without* ExSEnt (WoE), evaluated at *C*_rep_ with a fixed threshold of 0.5. For each cluster we report the number of misclassified subjects (*n*_err_) with FN/FP breakdown, the overlap (misclassified by both models), and the number uniquely misclassified by each model. The Jaccard index *J* = |WE ∩ WoE|*/*|WE ∪ WoE| summarizes how similar the error sets are (lower *J* indicates higher complementarity).

| Cluster | $N$ | WE $n_{\text{err}}$ (FN/FP) | WoE $n_{\text{err}}$ (FN/FP) | Overlap | WE-only | WoE-only | Union | $J$ |
| --- | --- | --- | --- | --- | --- | --- | --- | --- |
| LPFC | 45 | 14 (13/1) | 16 (13/3) | 9 | 5 | 7 | 21 | 0.43 |
| LVA | 55 | 15 (12/3) | 14 (9/5) | 11 | 4 | 3 | 18 | 0.61 |
| RPFC | 58 | 19 (15/4) | 20 (13/7) | 9 | 10 | 11 | 30 | 0.30 |
| RVA | 67 | 24 (16/8) | 21 (14/7) | 11 | 13 | 10 | 34 | 0.32 |

Taken together, these subject-level patterns support the interpretation that ExSEnt and non-ExSEnt features are complementary, as they capture different aspects of disease-related variability across subjects. This motivates late-fusion strategies (e.g., probability averaging or stacking WE and WoE predictors, optionally combined across regions), which can in principle reduce errors toward the intersection set (subjects misclassified by both) rather than the union, while preserving interpretability by tracking which feature family dominates each subject’s decision.

## 4. Discussion

Across the examined cortical clusters, the stability-selection analysis indicated that ExSEnt-derived features constitute a major driver of the learned decision function, with region- and hemisphere-dependent impact. Throughout the study, ExSEnt measures emerged as the most frequently selected and consistently weighted predictors across three of the four regions (LPFC and both VA), where the top-ranked features were from the ExSEnt family, with stability-selection frequencies exceeding 70% within training subsamples. This pattern indicated robust cross-subject reproducibility across spatially distinct sources. In the left hemisphere, this predominance translated into clear gains in discriminative performance: LPFC achieved the strongest classification when ExSEnt was included (AUC = 0.861, BAcc = 82.6%), whereas removing ExSEnt substantially degraded performance (Table 4). This drop was accompanied by a marked shift in the final stable signature (Table 5), where the ExSEnt-driven LPFC model was dominated by three perfectly stable theta/alpha ExSEnt features with large negative coefficients, consistent with a robust inverse association with dementia. In the LVA, the discriminative performance was high in both feature sets, indicating that ExSEnt contributed complementary information, but was not strictly necessary to preserve separability in this region. Thus, the left-hemisphere advantage was not uniform: ExSEnt for the LPFC yielded the best result, but it improved net performance only marginally in LVA, despite reshaping the selected biomarker composition toward *α*-band ExSEnt descriptors.

In contrast, right-hemisphere clusters remained closer to chance-level discrimination irrespective of feature set, and ExSEnt did not yield systematic improvements. In RPFC, performance was low for both conditions, with a slight advantage when ExSEnt was excluded; in both cases TAR^md^ was the most stable feature. Nevertheless, ExSEnt still contributed in RPFC in the sense of feature stability: ExSEnt metrics such as H*_A_* in the *α* band and H*_D_* in the *β* band ranked immediately below the dominant power-based terms and exhibited comparatively large coefficient magnitudes, suggesting a meaningful association with cognitive status that did not consistently improve held-out generalization under the current constraint, but performed better than other non-linear measures in this study. In RVA, excluding ExSEnt yielded modestly higher AUC and BAcc. The top feature in the WE model was a stable high-*γ* ExSEnt feature with a negative coefficient together with positive TAR and Hurst exponent.

The most stable ExSEnt measures in LPFC carried negative coefficients, indicating an inverse association with the probability of dementia: higher values of these features were linked to a lower likelihood of dementia. This pattern suggested that dementia cases exhibited a systematic reduction in *(i)* the temporal irregularity of local oscillatory timing (more predictable inter-extrema spacing), and *(ii)* the coupled timing–amplitude irregularity captured by H*_DA_*, alongside a reduced cross-window fluctuation of these irregularity measures.

This interpretation was consistent with the broader electrophysiological literature reporting that Alzheimer’s disease and prodromal stages are characterized not only by spectral slowing but also by a loss of dynamical complexity (lower entropy/irregularity), with particularly prominent effects in frontal regions at shorter temporal scales [96, 97]. Entropy-based analyses (e.g., SampEn/MSE and related measures) have repeatedly shown reduced complexity in AD relative to age-matched controls, and multiscale approaches have specifically reported reduced small-scale complexity in frontal areas together with links to cognitive status [98, 99]. A plausible mechanistic account is that early pathology disrupts the richness of prefrontal network dynamics through synaptic degradation and “cortical disconnection”, reducing the diversity of effective interactions; computational and empirical work supports a direct relationship between reduced functional connectivity and decreased EEG complexity [100]. Within this framework, the LPFC ExSEnt findings may be interpreted as an electrophysiological readout of diminished dynamical repertoire in executive-control networks: a shift toward more regular, less variable timing of local extrema and weaker timing–amplitude irregularity, consistent with reduced adaptive capacity of the underlying circuitry.

From a biomarker design perspective, these LPFC results motivate the use of ExSEnt-derived timing irregularity as an interpretable marker of early dementia-related network degradation. Because ℋ*_𝒟_* and ℋ*_𝒟𝒜_* explicitly quantify the entropy of inter-extremum timing (and its coupling to amplitude structure), they provide a mechanistically grounded complement to classical slowing indices (e.g., TAR): rather than tracking mean frequency shifts, they index the breakdown of temporal organization. In practice, these findings support compact LPFC-focused classifiers driven by a small set of ExSEnt predictors and motivate longitudinal analyses of whether LPFC ExSEnt timing irregularity tracks cognitive decline (e.g., MMSE) beyond normal aging.

The subject-level overlap analysis clarified how ExSEnt improves performance beyond aggregate metrics. In LPFC, the low Jaccard overlap and the concurrent reduction in false negatives indicated that ExSEnt contributed decision evidence that was non-redundant with the remaining feature families, rescuing a subset of dementia subjects that were otherwise missed. By contrast, in LVA and especially RPFC, the high overlap and near-threshold posterior probabilities suggested a small classification margin under the current LOSO setting, such that including ExSEnt primarily perturbed borderline cases rather than resolving a distinct error subset, consistent with limited net gains at a fixed threshold. The wider probability spread of ExSEnt-associated errors in RVA further indicated that ExSEnt reshaped the confidence structure of the classifier, not only the predicted labels, implying region-dependent effects on separability. Overall, these results motivate combining the two feature sets (and possibly the regions) to reduce errors on the subjects that both models find difficult. This keeps the model interpretable because we can still track which feature family drives each subject’s decision.

Several classification methods have been applied to this dataset in recent literature, with varying degrees of success. In the original study by Miltiadous et al. [77], a Random Forest classifier achieved an initial accuracy of 77.0%. Subsequent refinements reported higher binary accuracies, such as 83.06% for Alzheimer’s Disease versus Healthy Control using LightGBM and complexity-based features [101], and 83.28% using a hybrid Convolution-Transformer architecture known as DICE-net [102]. Recent attempts to improve discrimination have relied on even more intricate methods, such as quantifying phase-amplitude coupling and communication between electrode pairs, which achieved classification accuracies of 76.9% and, 90.4% for AD and FTD vs Healthy, respectively, but required extensive connectivity modeling [103]. However, these results typically rely on computationally intensive and lengthy analysis pipelines, such as those based on functional connectivity across the entire electrode montage or intricate signal decomposition. In contrast, the approach proposed in this study utilizes a significantly simpler analysis focused on only a single brain source, yet it achieves comparable state-of-the-art results with high interpretability.

## 5. Conclusion

Our findings confirm and extend the current understanding of neurophysiological alterations in early dementia in two complementary directions. First, the distribution of stable predictors across frequency bands indicated that although alpha–theta slowing remained a canonical signature of cognitive decline, when disentangled temporal and amplitude driven entropies were added to the feature set, this new information provided better discriminative values, making them the most frequently chosen features compared to the majority of the classic measures. ExSEnt measures from low to high frequency bands appeared as the most stable discriminative measures, implying that dementia-related changes affect cortical dynamics in a wide range of frequency bands, beyond shifts in mean oscillatory content. Moreover, the observed hemispheric asymmetry in performance and feature composition suggests lateralized vulnerability: left-hemisphere networks (LPFC and LVA) yield consistently higher discriminability than their right-hemisphere counterparts, compatible with earlier or stronger disruption of left-lateralized fronto-temporal circuits in the examined cohort.

Second, the sign of the coefficients helps the physiological interpretation of these spectral patterns. In LPFC, the dominant ExSEnt predictors had negative weights, showing that the dementia group is marked by lower temporal and amplitude–temporal coupling entropies in theta–alpha bands. Neurophysiologically, this is compatible with a shift toward less diverse prefrontal dynamics and reduced temporal flexibility of local network activity—information that is not captured by spectral slowing. By contrast, right-hemisphere models show weaker and less consistent separation. In the beta band, they had positive weights, suggesting relatively increased irregularity in the dementia group, while high gamma band time-amplitude entropies showed reduction with dementia. This pattern could reflect compensatory engagement, altered excitation–inhibition balance, or stage-dependent reorganization, indicating that early dementia may involve both reduced variability within vulnerable prefrontal networks and a redistribution of dynamical organization elsewhere.

In particular, even when classical entropy measures were included among the candidate predictors, ExSEnt emerged as the strongest discriminative family, producing stable and reproducible classification performance in the LPFC. This suggests that explicitly separating temporal irregularity from amplitude-driven irregularity provides a more informative representation in which pathology-related dynamical changes are expressed more clearly.

From a biomarker and translational perspective, the primary conclusion is that the ExSEnt family provides a robust, interpretable gain in EEG-based dementia discrimination, with its most pronounced benefit localized to LPFC. Across regions, ExSEnt measures are repeatedly selected with high cross-fold stability, demonstrating that the extrema-segmented entropy-based representation captures reproducible disease-related structure. Critically, in LPFC the inclusion of ExSEnt translates into a substantial performance improvement under strict cross-subject evaluation, and it yields compact decision rules dominated by a small set of highly stable ExSEnt predictors. In contrast, the right-hemisphere clusters remain closer to chance regardless of feature set, indicating that the dominant discriminative signal in this dataset is left-lateralized. Also, we would like to emphasize that the application of ExSEnt measures to the biosignals could enhance the classification and diagnostic precision when combined with imaging data from other modalities (i.e., MRI, fNIRS, etc.).

Finally, these results motivate two practical implications for early dementia detection through explainable-by-design AI. First, incorporating ExSEnt alongside established nonlinear and spectral descriptors can improve sensitivity to subtle preclinical alterations, particularly in prefrontal networks implicated in executive and memory-related control. Second, the subject-level error patterns indicate that ExSEnt and non-ExSEnt feature sets can be partially complementary, suggesting that principled fusion strategies (e.g., region-wise or feature-family-wise late fusion) may further reduce misclassification by leveraging distinct evidence streams while maintaining interpretability. Overall, by quantifying both stability and generalization under nested LOSO evaluation, this work supports ExSEnt as a promising, physiologically grounded biomarker family for early dementia screening from short resting-state EEG segments, with LPFC emerging as the most informative cortical target in the present cohort.

This dataset comprised 88 subjects and provides a solid foundation for method development; the observed effects are statistically robust, and the results are encouraging. Evaluating the ExSEnt family as a novel biomarker group in larger and more diverse cohorts will allow a more stringent assessment of generalizability and will further clarify their stability and reproducibility across populations, sites, and acquisition protocols.

## Code availability

All the codes used for preprocessing and analysis in this study are available at: /github.com/GNB-UAM.

## CRediT authorship contribution statement

S. Kamali: Conceptualization, Investigation, Software, Methodology, Formal analysis, Data curation, Visualization, Writing - original draft. F. Baroni: Conceptualization, Methodology, Validation, Resources, Writing – Review & Editing, Supervision. P. Varona: Conceptualization, Methodology, Validation, Resources, Writing – Review & Editing, Supervision, Project administration, Funding acquisition.

