## Supplementary tables for "ExSEnt for explainable dementia detection: disentangling temporal and amplitude-driven complexity boosts EEG-based classification"

### Supplementary Material

#### Supplementary Tables

**Table S1:** Table S1. LPFC misclassified subjects (with ExSEnt).

| $\text{ID}_{subject}$ | Group | Age | Gender | MMSE | True class | $\text{Class}_{pred}$ | $\text{Class}_{pedprob}$ | Error type |
|---|---|---|---|---|---|---|---|---|
| 2 | A | 78.0 | F | 22.0 | Case | Healthy | 0.027640 | FN |
| 68 | F | 78.0 | M | 25.0 | Case | Healthy | 0.041395 | FN |
| 29 | A | 53.0 | F | 16.0 | Case | Healthy | 0.087882 | FN |
| 9 | A | 77.0 | F | 23.0 | Case | Healthy | 0.220653 | FN |
| 86 | F | 49.0 | M | 26.0 | Case | Healthy | 0.427534 | FN |
| 7 | A | 79.0 | F | 20.0 | Case | Healthy | 0.468953 | FN |
| 54 | C | 78.0 | M | 30.0 | Healthy | Case | 0.742059 | FP |
| 44 | C | 64.0 | F | 30.0 | Healthy | Case | 0.583848 | FP |

**Table S2:** Table S2. LPFC misclassified subjects (without ExSEnt).

| $ID_{subject}$ | Group | Age | Gender | MMSE | True class | $Class_{pred}$ | $Class_{pedprob}$ | Error type |
|---|---|---|---|---|---|---|---|---|
| 11 | A | 71.0 | M | 22.0 | Case | Healthy | 0.124719 | FN |
| 67 | F | 66.0 | M | 24.0 | Case | Healthy | 0.187846 | FN |
| 74 | F | 53.0 | F | 20.0 | Case | Healthy | 0.224060 | FN |
| 68 | F | 78.0 | M | 25.0 | Case | Healthy | 0.235738 | FN |
| 29 | A | 53.0 | F | 16.0 | Case | Healthy | 0.237876 | FN |
| 9 | A | 77.0 | F | 23.0 | Case | Healthy | 0.267328 | FN |
| 66 | F | 73.0 | M | 20.0 | Case | Healthy | 0.280106 | FN |
| 16 | A | 68.0 | F | 14.0 | Case | Healthy | 0.313785 | FN |
| 88 | F | 55.0 | M | 24.0 | Case | Healthy | 0.488158 | FN |
| 73 | F | 57.0 | F | 22.0 | Case | Healthy | 0.495158 | FN |
| 43 | C | 72.0 | M | 30.0 | Healthy | Case | 0.996829 | FP |
| 38 | C | 62.0 | M | 30.0 | Healthy | Case | 0.837729 | FP |
| 50 | C | 68.0 | M | 30.0 | Healthy | Case | 0.685889 | FP |

**Table S3:** Table S3. LVA misclassified subjects (with ExSEnt).

| $ID_{subject}$ | Group | Age | Gender | MMSE | True class | $Class_{pred}$ | $Class_{pedprob}$ | Error type |
|---|---|---|---|---|---|---|---|---|
| 80 | F | 71.0 | F | 20.0 | Case | Healthy | 0.496944 | FN |
| 82 | F | 63.0 | M | 27.0 | Case | Healthy | 0.498721 | FN |
| 25 | A | 79.0 | F | 20.0 | Case | Healthy | 0.498831 | FN |
| 13 | A | 64.0 | F | 20.0 | Case | Healthy | 0.499044 | FN |
| 88 | F | 55.0 | M | 24.0 | Case | Healthy | 0.499118 | FN |
| 17 | A | 61.0 | F | 6.0 | Case | Healthy | 0.499186 | FN |
| 86 | F | 49.0 | M | 26.0 | Case | Healthy | 0.499292 | FN |
| 12 | A | 63.0 | M | 18.0 | Case | Healthy | 0.499401 | FN |
| 71 | F | 62.0 | M | 20.0 | Case | Healthy | 0.499469 | FN |
| 31 | A | 67.0 | F | 22.0 | Case | Healthy | 0.499591 | FN |
| 79 | F | 60.0 | F | 18.0 | Case | Healthy | 0.499730 | FN |
| 40 | C | 61.0 | M | 30.0 | Healthy | Case | 0.501313 | FP |
| 45 | C | 70.0 | F | 30.0 | Healthy | Case | 0.500365 | FP |

**Table S4:** Table S4. LVA misclassified subjects (without ExSEnt).

| $ID_{subject}$ | Group | Age | Gender | MMSE | True class | $Class_{pred}$ | $Class_{pedprob}$ | Error type |
|---|---|---|---|---|---|---|---|---|
| 80 | F | 71.0 | F | 20.0 | Case | Healthy | 0.497548 | FN |
| 13 | A | 64.0 | F | 20.0 | Case | Healthy | 0.498979 | FN |
| 31 | A | 67.0 | F | 22.0 | Case | Healthy | 0.499097 | FN |
| 12 | A | 63.0 | M | 18.0 | Case | Healthy | 0.499382 | FN |
| 82 | F | 63.0 | M | 27.0 | Case | Healthy | 0.499496 | FN |
| 18 | A | 73.0 | F | 23.0 | Case | Healthy | 0.499548 | FN |
| 79 | F | 60.0 | F | 18.0 | Case | Healthy | 0.499726 | FN |
| 25 | A | 79.0 | F | 20.0 | Case | Healthy | 0.499868 | FN |
| 40 | C | 61.0 | M | 30.0 | Healthy | Case | 0.500972 | FP |
| 63 | C | 66.0 | M | 30.0 | Healthy | Case | 0.500249 | FP |
| 45 | C | 70.0 | F | 30.0 | Healthy | Case | 0.500082 | FP |

**Table S5:** Table S5. RPFC misclassified subjects (with ExSEnt).

| $ID_{subject}$ | Group | Age | Gender | MMSE | True class | $Class_{pred}$ | $Class_{pedprob}$ | Error type |
|---|---|---|---|---|---|---|---|---|
| 6 | A | 61.0 | F | 14.0 | Case | Healthy | 0.499921 | FN |
| 67 | F | 66.0 | M | 24.0 | Case | Healthy | 0.499934 | FN |
| 14 | A | 77.0 | M | 14.0 | Case | Healthy | 0.499943 | FN |
| 18 | A | 73.0 | F | 23.0 | Case | Healthy | 0.499947 | FN |
| 73 | F | 57.0 | F | 22.0 | Case | Healthy | 0.499955 | FN |
| 25 | A | 79.0 | F | 20.0 | Case | Healthy | 0.499958 | FN |
| 28 | A | 49.0 | M | 20.0 | Case | Healthy | 0.499970 | FN |
| 33 | A | 72.0 | F | 20.0 | Case | Healthy | 0.499972 | FN |
| 80 | F | 71.0 | F | 20.0 | Case | Healthy | 0.499975 | FN |
| 16 | A | 68.0 | F | 14.0 | Case | Healthy | 0.499976 | FN |
| 2 | A | 78.0 | F | 22.0 | Case | Healthy | 0.499978 | FN |
| 10 | A | 69.0 | M | 20.0 | Case | Healthy | 0.499986 | FN |
| 12 | A | 63.0 | M | 18.0 | Case | Healthy | 0.499987 | FN |
| 9 | A | 77.0 | F | 23.0 | Case | Healthy | 0.499990 | FN |
| 66 | F | 73.0 | M | 20.0 | Case | Healthy | 0.499991 | FN |
| 78 | F | 62.0 | M | 22.0 | Case | Healthy | 0.499995 | FN |
| 74 | F | 53.0 | F | 20.0 | Case | Healthy | 0.499995 | FN |
| 83 | F | 68.0 | F | 20.0 | Case | Healthy | 0.499995 | FN |
| 22 | A | 68.0 | F | 20.0 | Case | Healthy | 0.499996 | FN |
| 40 | C | 61.0 | M | 30.0 | Healthy | Case | 0.500086 | FP |
| 55 | C | 67.0 | M | 30.0 | Healthy | Case | 0.500030 | FP |
| 60 | C | 71.0 | F | 30.0 | Healthy | Case | 0.500018 | FP |
| 47 | C | 70.0 | F | 30.0 | Healthy | Case | 0.500017 | FP |
| 38 | C | 62.0 | M | 30.0 | Healthy | Case | 0.500016 | FP |
| 57 | C | 64.0 | M | 30.0 | Healthy | Case | 0.500016 | FP |
| 49 | C | 62.0 | F | 30.0 | Healthy | Case | 0.500016 | FP |
| 46 | C | 63.0 | M | 30.0 | Healthy | Case | 0.500004 | FP |

**Table S6:** Table S6. RPFC misclassified subjects (without ExSEnt).

| $ID_{subject}$ | Group | Age | Gender | MMSE | True class | $Class_{pred}$ | $Class_{pedprob}$ | Error type |
|---|---|---|---|---|---|---|---|---|
| 66 | F | 73.0 | M | 20.0 | Case | Healthy | 0.499864 | FN |
| 67 | F | 66.0 | M | 24.0 | Case | Healthy | 0.499934 | FN |
| 18 | A | 73.0 | F | 23.0 | Case | Healthy | 0.499958 | FN |
| 73 | F | 57.0 | F | 22.0 | Case | Healthy | 0.499958 | FN |
| 16 | A | 68.0 | F | 14.0 | Case | Healthy | 0.499976 | FN |
| 2 | A | 78.0 | F | 22.0 | Case | Healthy | 0.499978 | FN |
| 6 | A | 61.0 | F | 14.0 | Case | Healthy | 0.499982 | FN |
| 25 | A | 79.0 | F | 20.0 | Case | Healthy | 0.499984 | FN |
| 10 | A | 69.0 | M | 20.0 | Case | Healthy | 0.499986 | FN |
| 12 | A | 63.0 | M | 18.0 | Case | Healthy | 0.499987 | FN |
| 9 | A | 77.0 | F | 23.0 | Case | Healthy | 0.499990 | FN |
| 80 | F | 71.0 | F | 20.0 | Case | Healthy | 0.499992 | FN |
| 24 | A | 69.0 | F | 20.0 | Case | Healthy | 0.499992 | FN |
| 14 | A | 77.0 | M | 14.0 | Case | Healthy | 0.499992 | FN |
| 33 | A | 72.0 | F | 20.0 | Case | Healthy | 0.499993 | FN |
| 78 | F | 62.0 | M | 22.0 | Case | Healthy | 0.499995 | FN |
| 74 | F | 53.0 | F | 20.0 | Case | Healthy | 0.499995 | FN |
| 83 | F | 68.0 | F | 20.0 | Case | Healthy | 0.499995 | FN |
| 40 | C | 61.0 | M | 30.0 | Healthy | Case | 0.500083 | FP |
| 63 | C | 66.0 | M | 30.0 | Healthy | Case | 0.500030 | FP |
| 52 | C | 73.0 | F | 30.0 | Healthy | Case | 0.500019 | FP |
| 60 | C | 71.0 | F | 30.0 | Healthy | Case | 0.500018 | FP |
| 38 | C | 62.0 | M | 30.0 | Healthy | Case | 0.500016 | FP |
| 49 | C | 62.0 | F | 30.0 | Healthy | Case | 0.500016 | FP |
| 47 | C | 70.0 | F | 30.0 | Healthy | Case | 0.500010 | FP |
| 57 | C | 64.0 | M | 30.0 | Healthy | Case | 0.500009 | FP |
| 46 | C | 63.0 | M | 30.0 | Healthy | Case | 0.500002 | FP |

**Table S7:** Table S7. RVA misclassified subjects (with ExSEnt).

| $ID_{subject}$ | Group | Age | Gender | MMSE | True class | $Class_{pred}$ | $Class_{pedprob}$ | Error type |
|---|---|---|---|---|---|---|---|---|
| 68 | F | 78.0 | M | 25.0 | Case | Healthy | 0.010549 | FN |
| 9 | A | 77.0 | F | 23.0 | Case | Healthy | 0.015104 | FN |
| 10 | A | 69.0 | M | 20.0 | Case | Healthy | 0.024198 | FN |
| 17 | A | 61.0 | F | 6.0 | Case | Healthy | 0.027878 | FN |
| 25 | A | 79.0 | F | 20.0 | Case | Healthy | 0.058028 | FN |
| 3 | A | 70.0 | M | 14.0 | Case | Healthy | 0.075169 | FN |
| 77 | F | 61.0 | M | 22.0 | Case | Healthy | 0.124952 | FN |
| 79 | F | 60.0 | F | 18.0 | Case | Healthy | 0.171513 | FN |
| 36 | A | 58.0 | F | 9.0 | Case | Healthy | 0.218131 | FN |
| 88 | F | 55.0 | M | 24.0 | Case | Healthy | 0.239480 | FN |
| 74 | F | 53.0 | F | 20.0 | Case | Healthy | 0.281751 | FN |
| 12 | A | 63.0 | M | 18.0 | Case | Healthy | 0.292407 | FN |
| 4 | A | 67.0 | F | 20.0 | Case | Healthy | 0.293796 | FN |
| 32 | A | 59.0 | F | 20.0 | Case | Healthy | 0.307009 | FN |
| 6 | A | 61.0 | F | 14.0 | Case | Healthy | 0.344582 | FN |
| 82 | F | 63.0 | M | 27.0 | Case | Healthy | 0.410872 | FN |
| 80 | F | 71.0 | F | 20.0 | Case | Healthy | 0.442869 | FN |
| 61 | C | 63.0 | F | 30.0 | Healthy | Case | 0.939750 | FP |
| 57 | C | 64.0 | M | 30.0 | Healthy | Case | 0.882013 | FP |
| 40 | C | 61.0 | M | 30.0 | Healthy | Case | 0.840625 | FP |
| 39 | C | 70.0 | M | 30.0 | Healthy | Case | 0.762207 | FP |
| 53 | C | 70.0 | M | 30.0 | Healthy | Case | 0.754994 | FP |
| 63 | C | 66.0 | M | 30.0 | Healthy | Case | 0.705861 | FP |
| 59 | C | 77.0 | M | 30.0 | Healthy | Case | 0.670591 | FP |
| 48 | C | 65.0 | M | 30.0 | Healthy | Case | 0.591386 | FP |
| 54 | C | 78.0 | M | 30.0 | Healthy | Case | 0.589913 | FP |
| 45 | C | 70.0 | F | 30.0 | Healthy | Case | 0.558642 | FP |

**Table S8:** Table S8. RVA misclassified subjects (without ExSEnt).

| $ID_{subject}$ | Group | Age | Gender | MMSE | True class | $Class_{pred}$ | $Class_{pedprob}$ | Error type |
|---|---|---|---|---|---|---|---|---|
| 10 | A | 69.0 | M | 20.0 | Case | Healthy | 0.494530 | FN |
| 3 | A | 70.0 | M | 14.0 | Case | Healthy | 0.498084 | FN |
| 20 | A | 71.0 | M | 4.0 | Case | Healthy | 0.498257 | FN |
| 9 | A | 77.0 | F | 23.0 | Case | Healthy | 0.498321 | FN |
| 77 | F | 61.0 | M | 22.0 | Case | Healthy | 0.498452 | FN |
| 68 | F | 78.0 | M | 25.0 | Case | Healthy | 0.498879 | FN |
| 82 | F | 63.0 | M | 27.0 | Case | Healthy | 0.499003 | FN |
| 25 | A | 79.0 | F | 20.0 | Case | Healthy | 0.499039 | FN |
| 29 | A | 53.0 | F | 16.0 | Case | Healthy | 0.499231 | FN |
| 18 | A | 73.0 | F | 23.0 | Case | Healthy | 0.499374 | FN |
| 74 | F | 53.0 | F | 20.0 | Case | Healthy | 0.499406 | FN |
| 71 | F | 62.0 | M | 20.0 | Case | Healthy | 0.499572 | FN |
| 12 | A | 63.0 | M | 18.0 | Case | Healthy | 0.499783 | FN |
| 36 | A | 58.0 | F | 9.0 | Case | Healthy | 0.499797 | FN |
| 79 | F | 60.0 | F | 18.0 | Case | Healthy | 0.499851 | FN |
| 32 | A | 59.0 | F | 20.0 | Case | Healthy | 0.499876 | FN |
| 88 | F | 55.0 | M | 24.0 | Case | Healthy | 0.499916 | FN |
| 6 | A | 61.0 | F | 14.0 | Case | Healthy | 0.499948 | FN |
| 59 | C | 77.0 | M | 30.0 | Healthy | Case | 0.501764 | FP |
| 45 | C | 70.0 | F | 30.0 | Healthy | Case | 0.501180 | FP |
| 40 | C | 61.0 | M | 30.0 | Healthy | Case | 0.500881 | FP |
| 61 | C | 63.0 | F | 30.0 | Healthy | Case | 0.500731 | FP |
| 43 | C | 72.0 | M | 30.0 | Healthy | Case | 0.500640 | FP |
| 38 | C | 62.0 | M | 30.0 | Healthy | Case | 0.500566 | FP |
| 60 | C | 71.0 | F | 30.0 | Healthy | Case | 0.500330 | FP |
| 53 | C | 70.0 | M | 30.0 | Healthy | Case | 0.500124 | FP |
